# Direct pharmacological AMPK activation inhibits mucosal SARS-CoV-2 infection by reducing lipid metabolism, restoring autophagy flux and the type I IFN response

**DOI:** 10.1101/2024.02.29.582713

**Authors:** Andrea Cottignies-Calamarte, Flora Marteau, Feifan He, Sandrine Belouzard, Jean Dubuisson, Daniela Tudor, Benoit Viollet, Morgane Bomsel

## Abstract

AMP-activated protein kinase (AMPK) plays a central role in regulating cell energy balance. When activated, AMPK supresses energy-consuming pathways such as lipid and protein synthesis while increasing nutrient availability through the activation of autophagy. These pathways downstream AMPK activation contribute to SARS-CoV-2 infection, which hijacks autophagy and accumulates lipid droplets in viral factories to support viral replication. Here, we assessed the antiviral activity of the direct pan-AMPK allosteric activator MK-8722 *in vitro.* MK-8722 efficiently inhibited infection of Alpha and Omicron SARS-CoV-2 variants in Vero76 and human bronchial epithelial Calu-3 cells at micromolar concentration. This inhibition relied on restoring the autophagic flux, which redirected newly synthesized viral proteins for degradation, and on a reduction in lipid metabolism, which affected the viral factories. Furthermore, MK-8722 treatment increased the type I interferon (IFN-I) response. Post-infection treatment with MK-8722 was enough to inhibit efficiently viral replication and restore the IFN-I response. Finally, MK-8722 treatment did not alter the SARS-CoV-2-specific CD8^+^ T cell response mounted upon Spike vaccination. Overall, by activating AMPK, MK-8722 acts as an effective antiviral against SARS-CoV-2 infection, even when applied post-exposure, paving the way for preclinical tests aimed at inhibiting viral replication and improving patients’ symptoms.

**Graphical abstract:** 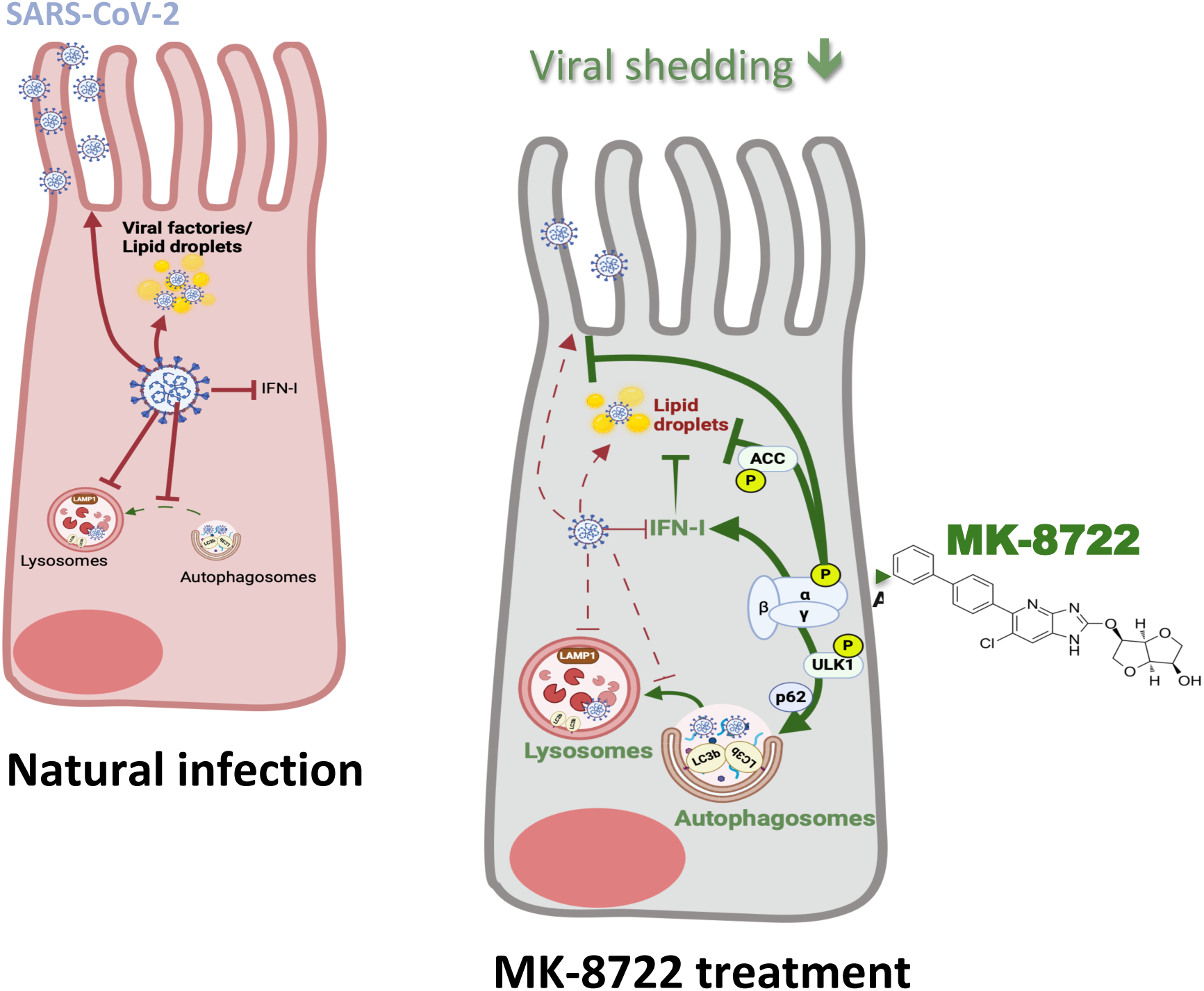

**Highlights:**

- MK-8722 exerts post-exposure antiviral activity
- MK-8722 induces a decrease in cellular lipid content
- MK-8722 promotes an increase in the autophagic flux of viral components
- MK-8722 promotes the restoration of the IFN-I activity
- MK-8722 antiviral activity is compatible with virus-specific T cell responses

## INTRODUCTION

The COVID-19 pandemic was efficiently contained due to the rapid development of vaccine and efficient public health policies. However, despite these efforts, SARS-CoV-2 Variants Of Concern (VOC), which are poorly sensitive to these vaccines, emerged and spread in the population, resulting in millions of deaths ^1^. Meanwhile, it has been challenging to develop an efficient antiviral treatment with an acceptable benefit/risk ratio. As of January 2024, the treatment of patients in the intensive care unit (ICU) remains primarily limited to drugs that either inhibit viral components such as Remdesivir and Paxlovid or target SARS-CoV-2-induced cytokine storm, such as the IL-6 receptor antagonist Tocilizumab ^1,2^.

After its nasal entry, SARS-CoV-2 primarily infects lung epithelial cells in a mechanism by which the viral envelope spike protein (Spike) binds to its main receptors angiotensin converting enzyme 2 (ACE2) and/or transmembrane protease serine 2 (TMPRSS2). Following viral entry and the delivery of viral RNA into the cytosol, the viral genes ORF1a and ORF1b are translated to the RNA-dependent RNA-polymerase (RdRp). The viral RdRp further generates the remaining viral mRNA and the viral cycle subsequently proceeds by hijacking cellular processes ^2^. Replication of SARS-CoV-2, similar to other enveloped RNA viruses, is supported by so-called viral factories with highly curved and three-dimensional folded membrane extensions of modified endoplasmic reticulum ^3,4^, which formation requires an increase in cell lipid metabolism. Viral factories compartmentalize viral genome from the cytosol for replication and transcription, and provide protection against host cell defences. Disrupting lipid metabolism hinders viral replication ^5–7^. Accordingly, SARS-CoV-2 infection results in a significant alteration in the cellular lipid repertoire in favor of polyunsaturated fatty acids, which are necessary for the generation of infectious viruses ^8–11^. This shift in lipid profile is mainly driven by the viral proteins ORF9, nsp4 and nsp6, leading to the accumulation of lipid droplets in infected cells *in vitro* and in COVID-19 autopsy tissues ^5,12–15^. Accordingly, disrupting chemically either of these mechanisms inhibits SARS-CoV-2 replication ^5,11,12,14,16^.

Throughout the SARS-CoV-2 viral cycle, autophagy is sequentially activated and inhibited ^17–20^. Indeed, autophagy is initiated by the early expressed nsp6, resulting in the formation of autophagosomes that are essential for the establishment of viral factories ^17,21^ and subsequent viral proteins expression ^17,21^. In turn, viral proteins OFR3a and ORF7a expressed at latter time post-infection prevent the fusion between autophagosomes and lysosomes thereby blocking completion of autophagy, as evidenced by increased LC3-B expression, activation of the ULK1 kinase and increase in the autophagy cargo receptor sequestosome-1/p62. Overall, this disruption of autophagy protects newly formed virus from degradation in the LAMP1^+^ lysosome ^19,22–25^.

SARS-CoV-2 infection also impairs the expression of Interferon type 1 (IFN-I) and Interferon Stimulated Genes (ISG), such as *Mx1* or *OAS1.* This reduction fails to restrict SARS-CoV-2 replication and this contributes to the hyperinflammatory symptoms observed in COVID-19 ^26–29^. Finally, SARS-CoV-2 infection interferes with the establishment of an efficient cellular response. By escaping lysosomal degradation, SARS-CoV-2 limits viral peptide loading onto MHC-I and subsequent presentation to specific cytotoxic CD8^+^ T Lymphocytes (CTL) ^30,31^. In addition, SARS-CoV-2 ORF8 down-regulates MHC-I expression from the plasma membrane, making infected cells less sensitive to SARS-CoV-2-specific CTLs ^32^.

AMP-activated protein kinase (AMPK) is a highly conserved cellular energy sensor that plays a crucial role in restoring the cellular energy balance by inhibiting ATP-consuming pathways (anabolism) and activating ATP-producing pathways (catabolism) ^33^. Notably, AMPK is a central regulator of lipid and protein metabolism. AMPK inhibits fatty acid and cholesterol synthesis, by direct phosphorylation in S79 and resulting inhibition of acetyl-CoA carboxylase (ACC) and 3-hydroxy-3-methylglutaryl-CoA (HMG-CoA) reductase ^33^. Additionally, AMPK is vital for the regulation of protein synthesis and cell growth through the inactivation of mTORC1 signalling through phosphorylation of Tuberous Sclerosis Complex 2 (TSC2) and Raptor at position S792 ^33^. Moreover, by activating Unc-51-like autophagy-activating kinase 1 (ULK1) after phosphorylating S555 and stimulating Nuclear factor erythroid 2-related factor 2 (NRF2)-dependent p62 increased expression, AMPK activation finely tunes cellular metabolic pathways ^33^. AMPK appears as a critical hub of cellular pathways affected by SARS-CoV-2 that are crucial for viral replication and pathogenesis.

A multitude of indirect and direct AMPK activators have been tested in preclinical models in various domains including viral infection and metabolic diseases ^33–35^. Recently, MK-8722, a direct allosteric-activator of AMPK, has been developed ^36^. MK-8722 is effective *in vitro* in the micromolar range on AMPK activation, compared to the millimolar range for metformin and 5-aminoimidazole-4-carboxamide riboside (AICAR) ^36,37^. MK-8722 has already been orally tested at 10 mg/kg in preclinical models of diabetes, resulting in improvements of glycemia but with a minor and reversible cardiac hypertrophy ^36^.

Here, we investigated whether allosteric AMPK activation by MK-8722 inhibits SARS-CoV-2 infection/replication by promoting autophagy of viral particles and restoring normal lipid metabolism and the IFN-I response.

## MATERIAL AND METHODS

### Cell culture and chemical compounds

All cells were cultured at 37°C in a humid atmosphere with 5% CO2. Vero76 (ATCC CRL-1587), Calu-3 (ATCC HTB-55) and Caco2 (ATCC HTB-37) cells were cultured in DMEM (Gibco 41966-029) containing 1% Penicillin-Streptomycin (PS) (Gibco 15140122) and decomplemented Foetal Calf Serum (FCS) (Eurobio CVFSVF06-01) at 10% (D10) for Calu-3 and Vero76 cells, and at 20% (D20) for Caco2 cells. Caco2 cells culture medium was supplemented with 1% MEM Non-Essential Amino Acids (Gibco 11140035). All viral inoculations and infections were made in DMEM containing 2% FCS (D2).

Peripheral blood mononucleated cells (PBMCs) of healthy donor were collected from the French blood bank (Etablissement Français du Sang, EFS) after initiation of SARS-CoV-2 vaccination in the global population (June 2021). PBMCs were obtained by Ficoll gradient centrifugation, aliquoted and stored in StemMACS CryoBrew (Miltenyi, 130-109-558) until use. CD8^+^ T cells were isolated from HLA-A2^+^ individuals using CD8 microbeads (Miltenyi 130-045-201) following manufacturer instructions. Cells (5×10^6^ cells at 10^6^ cells/mL) were further expanded with TransAct (Miltenyi, 130-111-160, 1:100 dilution) for 15 days in RPMI 10% FCS, 2 mM L-Glutamine (R10) (Thermo 25030081) + 10 IU/mL IL-2 (Roche 11011456001) after CD3/CD28 stimulation. TransAct was then washed out and cells frozen (1 million cells/vial) as described above. CD14-depleted PBMCs from healthy donors were obtained by collecting the flow-through of CD14 magnetic isolation (Miltenyi, 130-050-201). Resulting cells were then frozen as described above.

The absence of Mycoplasma in the cell lines was routinely controlled using the MycoAlert mycoplasma detection kit (Lonza, LT07-318).

Remdesevir and MK-8722 were purchased at MedChemExpress (respectively HY-104077 and HY-111363) and solubilized in DMSO at 10 µg/mL.

### Virus Propagation

SARS-CoV-2 variant Alpha and Omicron were obtained as we described ^38,39^, viral stocks prepared by amplification on Vero76 cells as we described ^35^,, aliquoted and cryopreserved at -80°C. Viral stocks were titrated on Vero76 cells after thawing as we described ^35^.

### Foci Forming Assay

Hundred thousand of Vero76 cells were seeded in 48 well-plates one day before infection, resulting in confluent cells at the day of infection. Virus-containing supernatant from infected cells to be tested were diluted at least 1,000 times in 100 µl of D2 and added to cells for 2h and then replaced with an overlay medium (300 µl) composed of 0.3% Low Melting Agarose (Sigma A0701) in D2 at 37°C. Plates were then placed at 4°C for 10 min to allow overlay medium to solidify and further incubated for 18h at 37°C. Plates were washed 3 times with PBS (Gibco 25300) to remove overlay, fixed with 4% paraformaldehyde (PFA) (Electron Microscopy Sciences 15710) in PBS (PBS-4% PFA) for 1h at room temperature and permeabilized with 0.1 % Triton-X100 in PBS for 10 min. Cells were washed 3 times and the viral nucleocapsid was labelled using an biotinylated-anti-N-antibody at 300 ng/mL (Bioss bsm-41411M-biotin) in PBS supplemented with 0.1% Saponin (Sigma S-7900), 2% FCS, and 2 mM EDTA (Perm Buffer) for 30 min at room temperature. Cells were washed three times in PBS, incubated with streptavidin-coupled-HRP 0.1 µg/mL (Vector Laboratories, INC, Burlingame, CA, 94010) in Perm Buffer for 30 min, washed again, and incubated with 100 µl/well of KPL TMB TrueBlue (Sera Care 5510-0030) for 30 min at 37°C.

The infectious index was established as the ratio of the viral copy number/mL over the FFU/mL titre computed from paired datasets obtained in RT-qPCR and titration experiments.

### Inhibition assay

Vero76 and Calu-3 cells were plated respectively at least 1 and 5 days before infection. Cells were treated with MK-8722 either continuously or only after infection (post-infection). Continuous treatment consisted in treatment at indicated concentrations for 1h before infection at 37°C, during the 2h inoculation, and until endpoint. Fully confluent cell layers were infected with an MOI of 0.001 for Vero76 cells or of 0.05 for Calu-3 in D2 for 2h. Inoculates were replaced by fresh D2 containing or not MK-8722, and cells further incubated for 1 and 4 days for Vero76 and Calu-3 cells, respectively. Before harvesting cells, supernatants were collected, clarified by centrifugation, and stored at -80°C for further RT-qPCR quantification and viral titration.

When indicated, untreated or pre-treated cells with 1 µM or 5 µM MK-8722 (for Vero76 and Calu-3 cells, respectively) were inoculated at 4°C for 1h to allow virus attachment to the cell membrane, wash three times in ice cold PBS and fresh D2, with or without MK-8722, was added before rising the temperature to 37°C for 1h or 24h for Vero76 and for 32h for Calu-3 cells, as indicated. Cells were then harvested and processed for *N* RNA quantification as described below. For post-infection treatments, MK-8722 was added at 8h post-infection (hpi) or 1 day post-infection (dpi), to reach indicated concentrations, controls receiving the same volume of D2.

### Quantification of infection by flow cytometry

Single-cell staining for viral protein and RNA by FISH-Flow was performed as described ^38^.

Briefly, cells were detached with trypsin 0.05% (Gibco 25300-024) for 10 min at 37°C and placed in 96 wells-plates. After centrifugation, cells were stained for 5min on ice for viability (Viobility 405/452, Miltenyi 130-130-420, diluted 100 times in cold PBS). Cells were washed in PBS and fixed with PBS-4% PFA. Cells were washed in Perm Buffer and labelled for 30min with anti-spike-AF488 antibody (R&D FAB105403G) diluted 250-fold in Perm Buffer and washed in Perm Buffer.

To further label viral RNA, cells were washed twice in Hybridization Wash Buffer (HWB) (2X SSC, 10% formamide (MP Biomedicals FORMD002), 0.2 mg/mL Bovine Seric Albumin (BSA) UltraPure (ThermoFisher Scientific AM2618) in ultrapure water (Invitrogen 10977-035)), resuspended in hybridization buffer (10% (w/v) Dextran sulfate (Calbiochem 265152), 1 mg Yeast tRNA (Invitrogen AM2616), 2X SSC, 10% Formamide, 0.2 mg/mL BSA RNase-free in ultrapure water) with 50 nM total SARS-CoV-2-quasar670-specific FISH probes (Stellaris) in equimolar proportion, targeting regions of S, ORF1 and N genes, resuspended in hybridization buffer and incubated for 12-16h at 37°C. After three washes in HWB, cells were resuspended in PBS and analysed by flow cytometry. Proportion of infected cells was quantified by flow cytometry (Guava 12-HT flow cytometer, Millipore) and analysed using the native GuavaSoft 3.4 software.

### Western-blotting

Cells were washed 3 times in PBS and lyzed with 1% Triton X-100 in 50 mM Tris pH 7.4, 150 mM NaCl, 1 mM EDTA, 1 mM EGTA, 10% Glycerol, 1 mM DTT, 1% Proteases Inhibitor Cocktail (Sigma P8340), 1% Phosphatases inhibitor cocktail II and III (Sigma P5725 et P0044 respectively). Cell lysates were clarified by centrifugation at 15,000g for 10min at 4°C, aliquoted and stored at -80°C. Protein content was quantified by BCA assay after sample thawing (Thermo Fischer J61522.AP). Around 25µg protein were separated by SDS-PAGE 12% at 100 mV and transferred on nitrocellulose membranes in wet condition at 80 mA at 4°C for 3h. Membranes were then saturated with TBS-0.5% Tween-20 (Sigma P1379, TBS-T)-5% BSA. After three washes in TBS-T, membranes were incubated with primary antibodies described in table 1 for 16h at 4°C. After three washes in TBS-T, relevant secondary antibody conjugated to HRP (see **Table 1**) was added for 1h at room temperature followed by washed 3 times in TBS-T. Finally, membranes were incubated with the West PicoPlus ECL HRP substrate (Thermo Fischer 34580). Signal was imaged with Fusion FX (Villber Lourmat) and quantified using Fiji ImageJ software. When indicated, data were computed from the Log2 fold change of the condition of interest over Non-infected non-treated cells and presented as heat map using GraphPad8.

**Table 1:** Antibody list used for western-blotting.

| **Table 1: Antibody list used for western-blotting** |  |  |  |  |
| --- | --- | --- | --- | --- |
| **Target** | **Host and antibody nature** | **Conjugation** | **Supplier & reference** | **Dilution** |
| **β-actin** | Goat polyclonal | - | Abcam ab8229 | 4,000 |
| **p-Thr172-AMPK** | Rabbit polyclonal | - | CST 2531 | 5,000 |
| **Pan-AMPK** | Rabbit polyclonal | - | CST 2532 | 5,000 |
| **p-Ser79-ACC** | Rabbit polyclonal | - | CST 11818 | 5,000 |
| **Pan-ACC** | Rabbit polyclonal | - | CST 3676S | 5,000 |
| **p-Ser555-ULK1** | Rabbit polyclonal | - | CST 5869 | 5,000 |
| **Pan-ULK1** | Rabbit polyclonal | - | CST 4773 | 5,000 |
| **p-Ser792-Raptor** | Rabbit polyclonal | - | CST 2083 | 5,000 |
| **Nucleocapsid** | Mouse IgG2b | Biotin | BIOSS bsm-41411M-biotin | 3,000 (300 ng/mL) |
| **Pan-Raptor** | Rabbit polyclonal | - | CST 2280 | 5,000 |
| **SQSTM1/p62** | Rabbit polyclonal | - | CST 5114 | 5,000 |
| **Goat** | Monkey polyclonal | HRP | Promega V805A | 5,000 |
| **Rabbit** | Donkey polyclonal | HRP | Southern Biotech 6440-05 | 5,000 |
| **Streptavidin** | - | HRP | Vector Laboratories, INC, Burlingame, CA, 94010 | 10,000 |

### RNA extraction, N normalizer plasmid and RT-qPCR

For RNA extraction, cells were washed 3 times in PBS and lysed in RA1 buffer. RNA was then extracted using Nucleospin RNA kit (Macherey-Nagel 740955) following manufacturer instructions, and stored at - 80°C before RT-qPCR experiments. Primers and probes, designed according to original CDC recommendations^40^, are described in **Table 2** (Eurofins Genomics). Normalizer plasmid was obtained after T-A cloning of the corresponding cDNA using High-Capacity RNA-to-cDNA kit (Thermo Fischer 387406) amplified using regular N-directed PCR with Platinium Taq (Thermo Fischer 10966034) following the TOPO cloning kit instruction (Thermo Fischer 10966034) and transformation of homemade thermocompetent TOP10 bacteria. Clones were selected using white-blue screening on LB agar plates with 100 µg/mL ampicillin and 20 µg/mL X-Gal (Thermo Fisher R0941). Plasmids were extracted using NucleoSpin Mini kit (Macherey Nagel 740588) and controlled by Sanger sequencing (Eurofins Genomics). For *Nucleocapsid* RNA relative or absolute quantification, RT-qPCR was performed using 4µl of cellular RNA and Taqman RNA-to-Ct 1step kit (Thermo Fischer 4392938) following manufacturer recommendations using a LightCycler 480 (Roche). For IFN-I and ISG expression analysis, reverse transcription was first carried out using High-Capacity RNA-to-cDNA and further quantified using Sybr Master mix I (Roche #04707516001). Relative gene expression over *β-actin* in cell lysates was calculated by the 2^-ΔΔCt^ method while absolute *N* abundance in supernatants was quantified using the N normalizer plasmid.

**Table 2:**
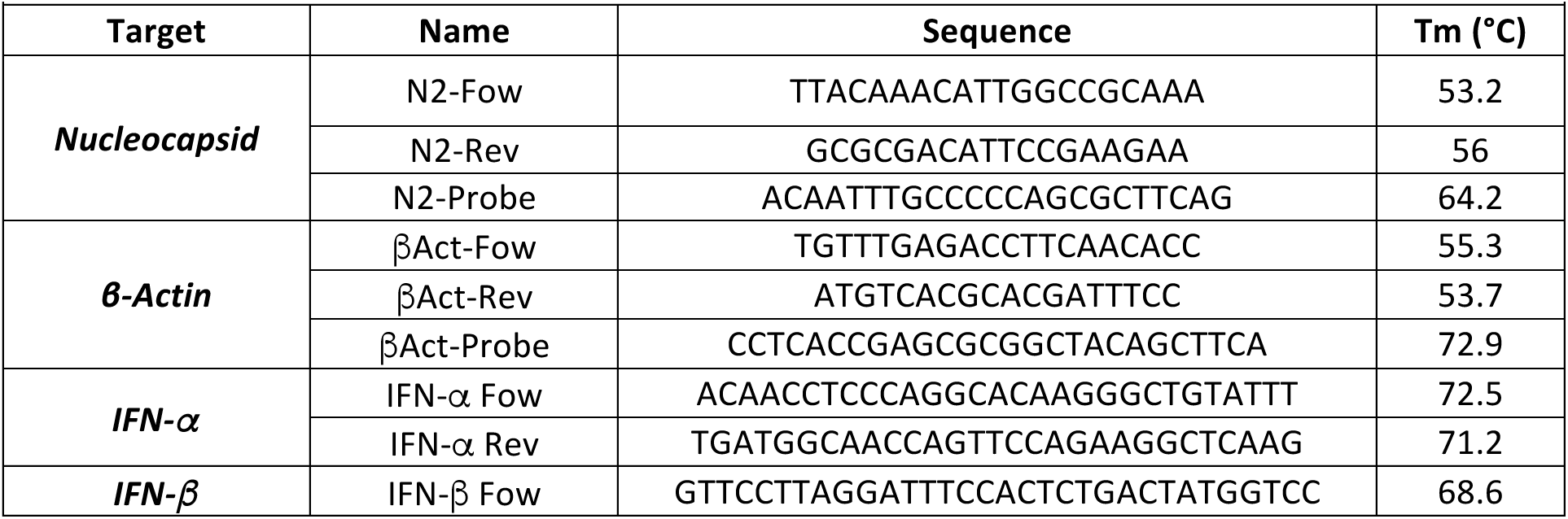
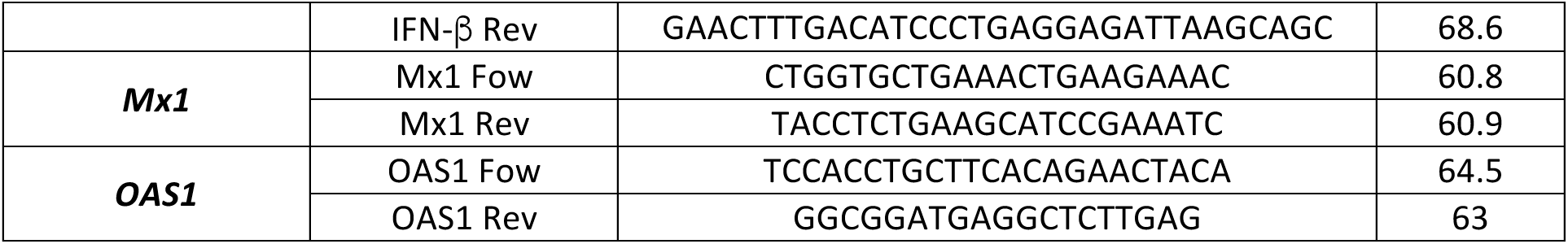
Primer characteristics.

### Immunofluorescence

Calu-3 cells were plated on Poly-L-Lysine (Sigma P8920) (0.1% in H_2_0) coated Ibidi slides (Ibidi 80841) at 1.2×10^5^/well for 5 days and infected with SARS-CoV-2 as described above. At 3dpi, cells were washed, stained with 10 µg/mL Nile Red (Sigma 72485) in PBS at 37°C for 20 min, and washed prior fixation, when indicated. Cells were fixed in PBS-4% PFA for 1h, washed and permeabilized using 0.1% TritonX-100 in PBS for 10 min. Non-specific binding was blocked with 5% FCS in PBS for 30 min. Cells were further labelled in Perm Buffer with biotinylated-anti-N antibody at 300 ng/ml (BIOSS bsm-41411M-biotin) and/or anti-LAMP1 at 625 ng/mL (BD 555798), or anti-LC3b antibody at 1 µg/mL (Novus Biological, #NB600-1384SS) and anti-LAMP1 antibody at 625 ng/mL, overnight at 4°C and the relevant secondary for 1h at room temperature (Streptavidin-AF488 Invitrogen S11223 at 2 µg/mL final, anti-mouse IgG1-Cy5 Abcam ab136127 at 1.5 µg/mL final or anti-Rabbit-Cy3 IgG at 1 µg/mL, Jackson 711-166-152, and anti-mouse IgG1-Cy5 at 1.5 µg/mL). After nuclear staining with DAPI, cells were washed, and slides mounted with Ibidi mounting medium (Ibidi 50001). Staining was observed by confocal microscopy (Xplore confocal microscope). At least five randomly chosen fields per condition were imaged and further analysed using ImageJ software and the particle analysis function. Lipid droplets (LD) were identified as Nile Red positive vesicular objects with diameter > 500 nm. Lysosomes were identified as LAMP1 positive vesicular objects with diameter > 700 nm. Analyse of lysosomes and LD was performed using Z-projections made from 6 consecutive slices 300 nm apart. The number of particles per infected cells, their mean area and intensity were then calculated. Colocalization on z-stack images was analysed with the JACoP plugin using object-based colocalization and setting threshold at 137, 134 and 120 for Lysosomes, LD and nucleocapsid objects, respectively. Manders’ Overlap Coefficients (MOC) corresponding respectively to green objects colocalizing in red objects and inversely red objects colocalizing in green objects were computed.

### Lysosomal function analysis

Confluent Calu-3 cells in a 96-well plate were inoculated with either Alpha or Omicron variant at 0.05 MOI for 2h, and cells were further cultured with fresh D2 up to 24hpi. MK-8722 at indicated concentration or D2 only were added for another 24h. Lysosomes were then stained with 5 nM Lysotracker RED DND-99 (ThermoFischer L7528) in DMEM for 30 min following the manufacturer instructions. Cells were then detached with trypsin 0.05% for 10 min, transferred into round bottom 96well-plate, stained for viability and fixed with PBS-4% PFA. After 20 min, cells were washed and the mean intensity of Lysotracker staining was quantified by flow cytometry as described above.

### Evaluation of MK-8722 treatment on T cell activation

CD14-depleted PBMCs were thawed and cultured overnight at 37°C in R10 supplemented with 10 UI/mL IL-2 and 1 µg/mL DNAse (StemCell, 07900). Cells were then washed, resuspend cells in R10 and distributed at 10^5^ cells/well in a round bottom 96-well plate. Transact stimulation, which is a combination of activating anti-CD3 and anti-CD28 antibodies (Miltenyi, 130-111-160), consisted in 100-fold final dilution of Transact in R10. Alternatively, Spike or Nucleocapsid Peptivator, T cell epitope peptide pools from either Spike or Nucleocapsid of SARS-CoV-2, respectively (Miltenyi 130-126-700 and 130-126-698, respectively) was used as stimulation source at a final concentration of 500 ng/mL in R10. After stimulation for 30 min, Brefeldin A (5 µg/mL final) containing anti-CD107a-PE-Cy5 antibody (BD 555802) following the manufacturer instruction was added and cells were incubated for 6h. Cells were then washed and stained with anti-CD3-APC-H7 (BD 560176) and anti-CD4-APC (BD 551980) following manufacturer instructions for 10 min on ice, followed by viability staining as described above and fixation in PBS-4% PFA. Cells were further stained overnight with anti-IFNγ-PE (Miltenyi, 130-113-498) following manufacturer instructions in Perm Buffer. Cells were washed in Perm Buffer and then in PBS. Frequencies of CD4^+^ and CD8^+^ T cells expressing CD107a^+^ or IFNγ^+^ were quantified by flow cytometry as described above. For each donor, we determined the net activation (NA) level for each marker after stimulation with Peptivator (NA(Peptivator)) by normalizing CD107a and IFNγ frequencies measured after stimulation with Peptivator over those measured after stimulation with Transact. This activation index reflects the capacity of the donor cells to respond to peptide-specific stimulation relative to its intrinsic stimulatory capacities.

### Evaluation of combined antiviral effects of MK-8722 and CD8^+^ T cells from vaccinated individuals on epithelial cell infection

CD8^+^ T cell activation being restricted to MHC-I, antigenic presentation occurs only between as antigen presenting cell (i.e. infected cell) and the T cells with similar HLA. We first showed that Caco2 cells, unlike Calu-3 cells, express the MHC-I molecule HLA-A2 and were thus capable of presenting antigens to HLA-A2-restricted CD8+ T cells from vaccinated individuals. Therefore, confluent Caco2 cells were inoculated for 1h with SARS-CoV-2 and further incubated for 8h in the absence of virus. Previously amplified CD8^+^ T cells were thawed in R10 complemented with 5IU/mL IL-2 to promote T cell survival, washed and added to Caco2 cells at 8hpi with or without 2.5 or 5 µM MK-8722. The co-cultures were further incubated at 37°C for 36h. T cells were then collected, stained with anti-CD3-APC-H7 (BD 560176) and anti-CD8-APC (BD 345775) following manufacturer instructions, and for viability before fixation as above. Cells were then intracellularly stained with anti-spike-AF488 (R&D FAB105403G) for 1h at room temperature. Infection was monitored in CD3^-^CD8^-^ cells by flow cytometry as described above.

### Graphical representation and statistics

Data and graphs were computed using Graphpad Prism (San Diego, V8). If variables followed a normal distribution after Shapiro-Wilk, *t*-test or ANOVA was calculated; otherwise, statistical analyses were performed using the Mann-Whitney or Kruskal-Wallis test. *p*-values <0.05 were considered significant. Tests were paired when experience design permitted.

## RESULTS

### MK-8722 inhibits SARS-CoV-2 replication in epithelial cells

We first evaluated the activity of the pan-AMPK allosteric activator MK-8722 in Vero76 cells. As early as 1h after treatment, MK-7288 activated AMPK and triggered downstream phosphorylation of ACC in a concentration dependent manner from 0.01 to 1 µM (**Fig. S1**). The antiviral activity of MK-8722 was then monitored in Vero76 cells infected by SARS-CoV-2 as schematized (**Fig. 1A**). The drug was added at 0.01 to 1 µM to the cells 1h before infection, during the inoculation with SARS-CoV-2 Alpha for 2h, and during the subsequent 24h chase. At 1dpi, Spike expression was quantified by flow cytometry (see gating strategy **Fig. S2A**). Spike^+^ cell frequency decreased in the presence of MK-8722, from 11.8±0.7% in untreated to 5±0.4% in 1 µM-treated conditions (ANOVA: p<0.001, **Fig. S2B**), corresponding to an inhibition of infection of 53±3% at 1 µM (ANOVA: p<0.0001, **Fig. 1B**). MK-8722 antiviral activity was further analysed in bronchial Calu-3 cells (See gating strategy **Fig. S2C**), which are more relevant to SARS-CoV-2 infection since they express TMPRSS2 and are competent for IFN-I signalling similar to bronchial cells *in vivo* unlike Vero76 cells. Infection of Calu-3 cells with alpha variant in the presence of MK-8722 significantly decreased the frequency of viral Spike^+^ and viral RNA^+^ cells in a dose-dependent manner compared to untreated cells (**Fig. S2D**). At 10 µM MK-8722, the frequency of Spike^+^ and viral RNA^+^ cells decreased compared to untreated cells, from 30.8±2.5% to 3.6±0.2%, and from 15.1±1% to 1.5±0.2%, respectively (ANOVA: *p*<0.0001, **Fig. S2D**). MK-8722 similarly inhibited Omicron infection, the frequency of Spike^+^ and viral RNA^+^ cells decreasing compared to untreated cells (compare 27.5±3% versus 4.5±0.6% Spike+ cell frequency and 31±3% vs 4.5±0.4% RNA+ cell frequency at 10 µM and without MK-8722, respectively) (**Fig. S2E**). These reductions were comparable to that induced by treating Calu-3 cells with the RNA-dependent RNA polymerase inhibitor Remdesivir at 1 µM (**Fig. S2F**). The IC50 of MK-8722 in Calu-3 cells against Alpha variant was reached at 0.7 µM and an IC90 at 9 µM (*t*-test: *p*<0.05, **Fig. 1C**). For the Omicron variant, MK-8827 showed an IC50 at 1,6 µM and an IC90 slightly above 10 µM (ANOVA: *p*<0.05, **Fig. 1D**). Both RNA^+^ and Spike^+^ frequencies decreased after MK-8722 treatment. As MK-8722 antiviral activity against Alpha and Omicron variants is similar, the drug likely targets the same pathways in the two cases. Furthermore, MK-8722 blocked viral production by Calu-3 cells in a concentration-dependent manner, as quantified by the decrease in *N* RNA by RT-qPCR from 1.5±0.14×10^9^ to 6±1.4×10^7^ viral copies/mL at 10 µM MK-8722 (ANOVA: *p*<0.0001) (**Fig. 1E**). MK-8722 did not alter the expression of ACE2 in Vero76 and Calu-3 cells, as measured by flow cytometry (**Fig. S2G**), nor impacted cell viability (**Fig. S2H**), indicative of a post-entry antiviral activity. Furthermore, we also investigated MK-8722 toxicity at 4h, 24h and 96h (Fig. S2I) and found that the drug was not toxic up to 10µM. The 50% toxicity dose was calculated to be 57µM at 96h in Calu3 cells. This allows us to determine a therapeutic index of 76 against the SARS-CoV-2 Alpha variant and 36 against the Omicron variant, thus, placing MK-8722 as an attractive antiviral candidate.

**Figure 1:**
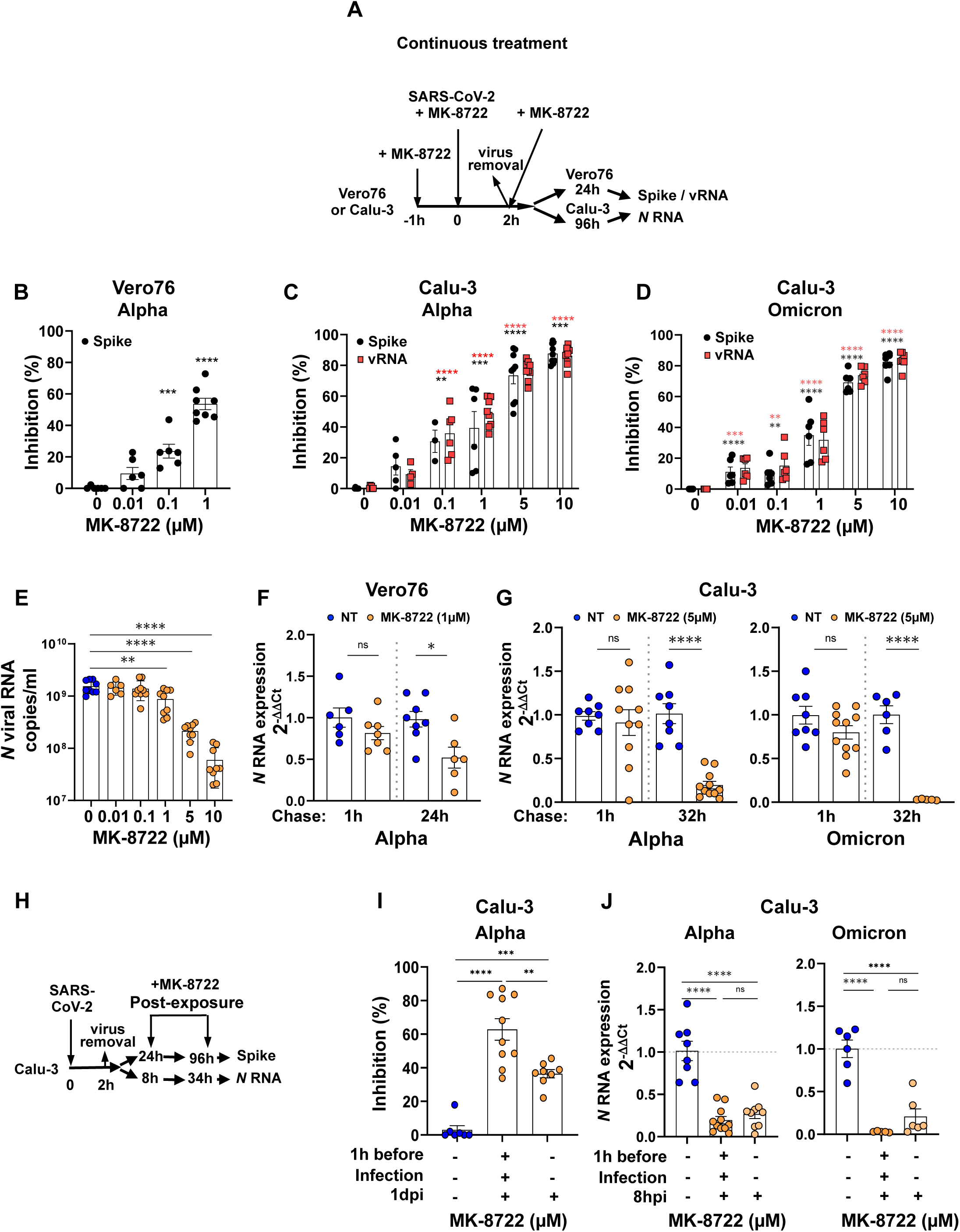
MK-8722 inhibits SARS-CoV-2 replication *in vitro*. **A:** ***Schematic outline of the continuous treatment protocol with MK-8722**.* **B-D: *Dose dependent inhibition of Vero76 and Calu-3 cell infection with SARS-CoV2 by continuous treatment with MK-8722***. Inhibition of infection with indicated SARS-CoV-2 variant in Vero76 was measured by flow cytometry (**B**) and that of Calu-3 by FISH-flow (**C-D**). n≥5 independent experiments. Labelling for SARS-CoV-2 Spike (Black) and for viral RNA (Red). **E: *Dose-dependent inhibition of viral genome release from Calu-3 cells*** infected with the Alpha variant after continuous treatment with MK-8722 measured by quantification of *N* RNA in the supernatant by RT-qPCR. n≥5 independent experiments. **F-G: *Inhibition of viral replication after MK-8722 treatment.*** Cells were incubated with the indicated SARS-CoV-2 variant in the absence or the presence of the drug (1 µM for Vero76 **(F)** and 5µM for Calu-3 cells **(G)**) for 1h at 4°C to synchronize infection and chased for 1h or for an additional 24h in the case of Vero76 cells and for 32h in the case of Calu-3 cells at 37°C. *N* RNA was quantified in cellular extracts by RT-qPCR. n≥5 independent experiments. ***H: Schematic representation of the post-exposure treatment protocol with MK-8722.*** **I-J**: ***Post-exposure treatment with MK-8722 is sufficient to block infection by SARS-CoV-2.*** Cells were left untreated or treated with MK-8722 (5 µM) either continuously with Alpha variant or treated post-infection as in **G**. Infection was evaluated by quantification of Spike expression by flow cytometry (**I**) or by RT-qPCR using the 2^-ΔΔCt^ method (**J**). Values represent infection inhibition. n≥6 independent experiments. Shown are mean ± SEM. ANOVA: * *p*<0.05, ** *p*<0.01, *** *p*<0.001, **** *p*<0.0001.

We next investigated whether MK-8722 exerts its antiviral effect at the early infection steps. After MK-8722 pre-treatment of Vero76 and Calu-3 cells (at 1µM and 5µM, respectively), cells were inoculated with Alpha or Omicron variants for 1h at 4°C to allow virus attachment. After virus removal, cells were incubated in the presence of the drug at 37°C to synchronize infection. Viral *N* RNA was quantified by RT-qPCR in the cells after a short (1h) or long (24h in Vero76 and 32h in Calu-3 cells) chase times. As control, cells were similarly infected but left untreated. Viral RNA accumulated after 1h chase in the two cell types for the two variants similarly in treated and non-treated conditions, although the drug had tendency to decrease relative *N* RNA levels in a non-significant statistical manner (1±0.12 vs 0.82±0.08 for Vero76, 1±0.05 vs 0.91±0.15 and 1±0.1 vs 0.8±0.08 for Alpha and Omicron variants in Calu-3 cells, respectively, *t-*test: ns in all conditions, **Fig. 1F** and **1G**). In contrast and as expected, *N* RNA expression was significantly reduced after a longer chase (1±0.09 vs 0.52±0.12 for Vero76, 1±0.11 vs 0.2±0.04 and 1±0.1 vs 0.03±0.003 for Alpha and Omicron variants in Calu-3 cells, respectively, *t*-test *p*<0.05, <0.0001 and <0.0001, respectively). These results suggest that MK-8722 has little antiviral effect during the early steps of infection, namely adsorption and fusion, but acts in a mechanism controlling post-entry replication steps.

### Post-infection treatment with MK-8722 is sufficient to inhibit SARS-CoV-2 replication

We next evaluated if MK-8722 was also active on cells after infection (**Fig. 1H**). Post-exposure treatment at 1dpi and until harvest at 3dpi efficiently blocked viral replication, quantified by Spike expression, by 36±7% (**Fig. 1I**, ANOVA: *p*<0.0001), although still less than the continuous treatment, which reduced infection by 63±6% (ANOVA: *p*<0.0001). Continuous and post-exposure treatment with MK-8722 (5 µM) resulted in a similar inhibition of viral genomes accumulation of both Omicron and Alpha variants in infected cells (**Fig. 1J**). Indeed, after continuous treatment with MK-8722 for 34h, *N* RNA relative amount decreased significantly by 80±0.04% and by 97±0.7% in Alpha and Omicron infected cells, respectively (*t*-test: *p*<0.0001 for both variants vs untreated cells). Limiting the MK-8722 treatment to the post-exposure phase, namely from 8hpi, decreased *N* RNA relative amount by 75±0.05% and 80±20% in Alpha and Omicron infected cells, respectively (ANOVA: *p*<0.0001 for both Alpha and Omicron variants vs untreated cells). Altogether, these data suggest that post-exposure treatment by MK-8722 is sufficient to inhibit viral infection.

### MK-8722-induced AMPK activation triggers autophagy induction and inhibits lipids metabolism to control SARS-CoV-2 replication

We further examined the mechanism of SARS-CoV-2 inhibition by MK-8722. Therefore, the modification by the drug of the signalling cascade downstream AMPK activation during SARS-CoV-2 infection of Vero76 and Calu-3 cells relative to no treatment was analysed using western blot (**Fig. S3A-B**). Quantification of the western blots relative to non-infected non-treated control condition is shown in **Fig. 2A** and summarized in a heat map in which values were expressed as log2 fold change (Log2(FC)) (**Fig. 2B**). As expected, MK-8722 treatment of both Vero76 and Calu-3 cells significantly activated AMPK that became phosphorylated at position S172 (0.85±0.4 and 1.2±0.3 log2(FC), t-test: p<0.05 and p<0.01, respectively, **Fig. 2B**). AMPK phosphorylation was comparable in infected and non-infected cells (0.0±0.5 and -0.02±0.3 log2(FC), respectively), while MK-8722 treatment stimulated AMPK phosphorylation even more in infected than in non-infected cells (3.1±0.4 and 2.3±0.9 log2(FC), respectively, t-test: p<0.05 both). As expected, the drug blocked infection as shown by a significant decrease in N protein levels in both Vero76 and Calu-3 cells (-0.88±0.2 and -1.7±0.4 Log2(FC), respectively, t-test: p<0.05 both, **Fig. 2A-B**).

**Figure 2:**
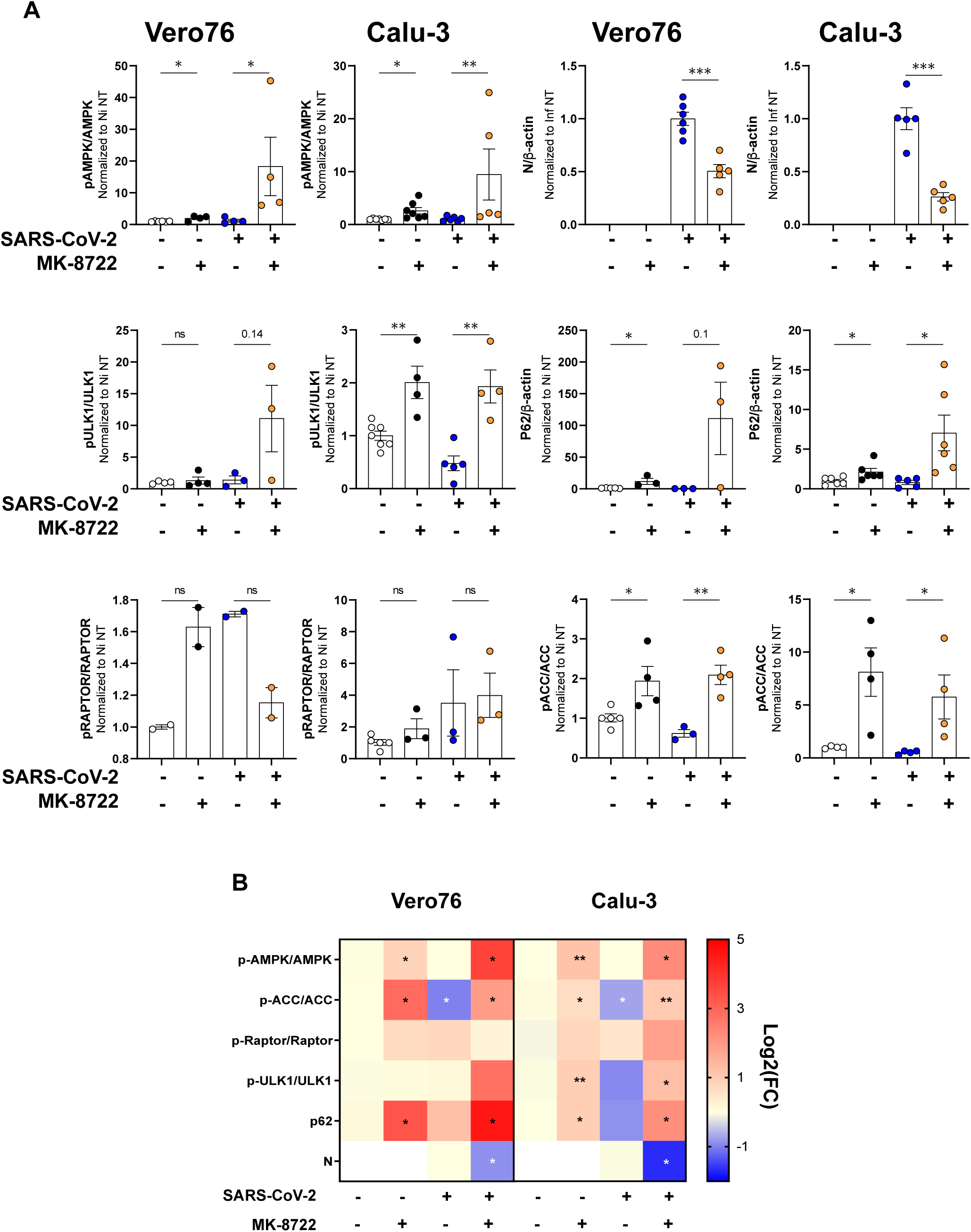
MK-8722-dependent AMPK activation inhibits SARS-CoV-2 replication by inhibition of the lipid biosynthetic pathway and restoration of the autophagy flux. **A**: **Activation or expression ratio of AMPK, ACC, raptor, ULK1, p62 and Nucleocapsid were analyzed by western blot** in Vero76 and Calu-3 cells after continuous MK-8722 treatment as indicated in Figure 1A Western blots were quantified by ImageJ, and each marker value in indicated condition was normalized over corresponding marker expression value in the non-infected non-treated condition. Shown are mean ± SEM. n≥2 independent experiments. **B: *Quantification of the western blot presented as Heat maps:*** Heat maps represent the transformation by Log2 of the expression/phosphorylation values quantified on each blot by ImageJ relative to corresponding non-infected non-treated condition for Vero76 and Calu-3 cells. n≥2 independent experiments. Student’s *t*–test: * *p*<0.05, ** *p*<0.01.

We then monitored S555 ULK1 phosphorylation and p62 relative expression as markers of the initiation of autophagy, Raptor phosphorylation as a marker of protein synthesis inhibition, and S79 ACC phosphorylation as a marker of the disruption of lipid metabolism. While neither MK-8722 nor SARS-CoV-2 infection alone affected ULK1 phosphorylation (with the exception of Calu-3 cells, non-infected but treated, where increase reached 1±0.2 Log2(FC), *p<*0.01), MK-8722 treatment during infection was associated with an increase in ULK1 S555 phosphorylation in both cell types (from 0.05±0.5 to 2.8±1.2 Log2(FC) in Vero76 cells and from -0.9±0.5 to 1.3 ±0.6 Log2(FC) in Calu-3 cells, t-test: ns and p<0.05, respectively, **Fig. 2A-B**). Concomitantly, p62 expression strongly and significantly increased in both cell types (3.33±0.56 and 2±0.6 Log2(FC) for Vero76 and Calu-3 cells, respectively, t-test p<0.05, **Fig. 2A-B**). S792 Raptor phosphorylation remained unchanged whatever the conditions. Finally, SARS-CoV-2 infection strongly decreased S79 ACC phosphorylation significantly (-1±0.29 and -0.7±0.2 Log2(FC), respectively and *t*-test: *p<*0.05, **Fig. 2A-B**) while it was significantly increased by MK-8722 treatment during infection (1.9±0.5 and 1±0.2 Log2(FC), *t*-test: *p*<0.05 and *p*<0.01, respectively, **Fig. 2A-B**).

Altogether these results suggest that the antiviral mechanism of MK-8722 relies on the AMPK activation-dependent restoration of the autophagic flux that is otherwise blocked by SARS-CoV-2 proteins, due to ULK1. Phosphorylation of ULK1 and increased p62 expression likely result in an increased access of viral components to phagosomes and further phagosomes fusion with lysosomes, resulting in subsequent degradation of viral proteins in phagolysosomes, preventing viral dissemination. As complementary anti-viral mechanism, MK-8722 treatment increases ACC phosphorylation that decreases lipid synthesis, which may reduce lipid droplets content and thus affect the establishment of viral factories. However, as shown by the lack of Raptor phosphorylation, the antiviral activity of MK-8722 does not involve a shutdown of viral protein synthesis. Trends in all marker changes after MK-8722 treatment were similar between cell lines indicating a common antiviral mechanism. We therefore focused our study on Calu-3 cells since they are more relevant to SARS-CoV-2 infection.

### MK-8722 treatment perturbs the viral factory establishment

We next explored the capacity of MK-8722 to perturb the distribution of lipids associated with viral factories and lipid droplets in cells infected with the Alpha variant. Therefore, lipid reorganisation in infected Calu-3 cells was monitored after continuous treatment with MK-8722 by staining lipids with the lipophilic marker Nile Red analysed by confocal microscopy. As expected ^34^, treatment with MK-8722 in uninfected cells blocked almost entirely the synthesis of lipids, as shown by the loss of Nile Red staining (**Fig. S4A**). Infection increased the overall Nile Red staining by 40±10% compared to non-infected cells (**Fig. S4B**, ANOVA: *p<*0.05), but also dramatically modified Nile Red cellular distribution into aggregates likely corresponding to lipid droplets (compare **Fig. 3A, S4B**). This overall infection-induced increase in Nile Red staining was abolished by MK-8722 treatment (**Fig. S4B**, 0.943±0.14-fold change, infected non-treated vs treated cells, ANOVA: *p<*0.05) and returned to untreated uninfected cell level. Furthermore, in infected cell, identified by viral N protein labelling, MK-8722 significantly reduced the number, area and intensity of lipids droplets (as defined in the method section) (**Fig. 3A** and **3B**, 0.29±0.09 and 0.28±0.1 and 0.33±0.15 fold-change over infected non-treated cells, *t*-test: *p<*0.01, *p<*0.05 and *p<*0.05, respectively). We then evaluated if the N protein, a maker of virus replication in viral factories, was redistributed out of lipid droplets by the drug. In the absence of MK-treatment, N protein and lipids colocalized as expected, although partially, with Manders’ Overlap Coefficient (MOC) being in the range of previous reports ^12^. This colocalisation decreased significantly in cells infected in the presence of MK-8722, as indicated by lower MOCs for N protein in Nile Red (0.085±0.024 vs 0.01±0.005, Mann-Whitney *p<*0.001) and for Nile Red in N protein (0.16±0.05 vs 0.027±0.01, Mann-Whitney *p<*0.01) in MK-8722 treated vs untreated infected cells (**Fig. 3A** and **3C**). Altogether, SARS-CoV-2 infection increases the accumulation of lipid droplets with which viral components colocalise. MK-8722 treatment reverses these effects, likely reflecting an alteration of cell lipid composition following increased ACC phosphorylation, thus limiting the production of viral factories.

**Figure 3:**
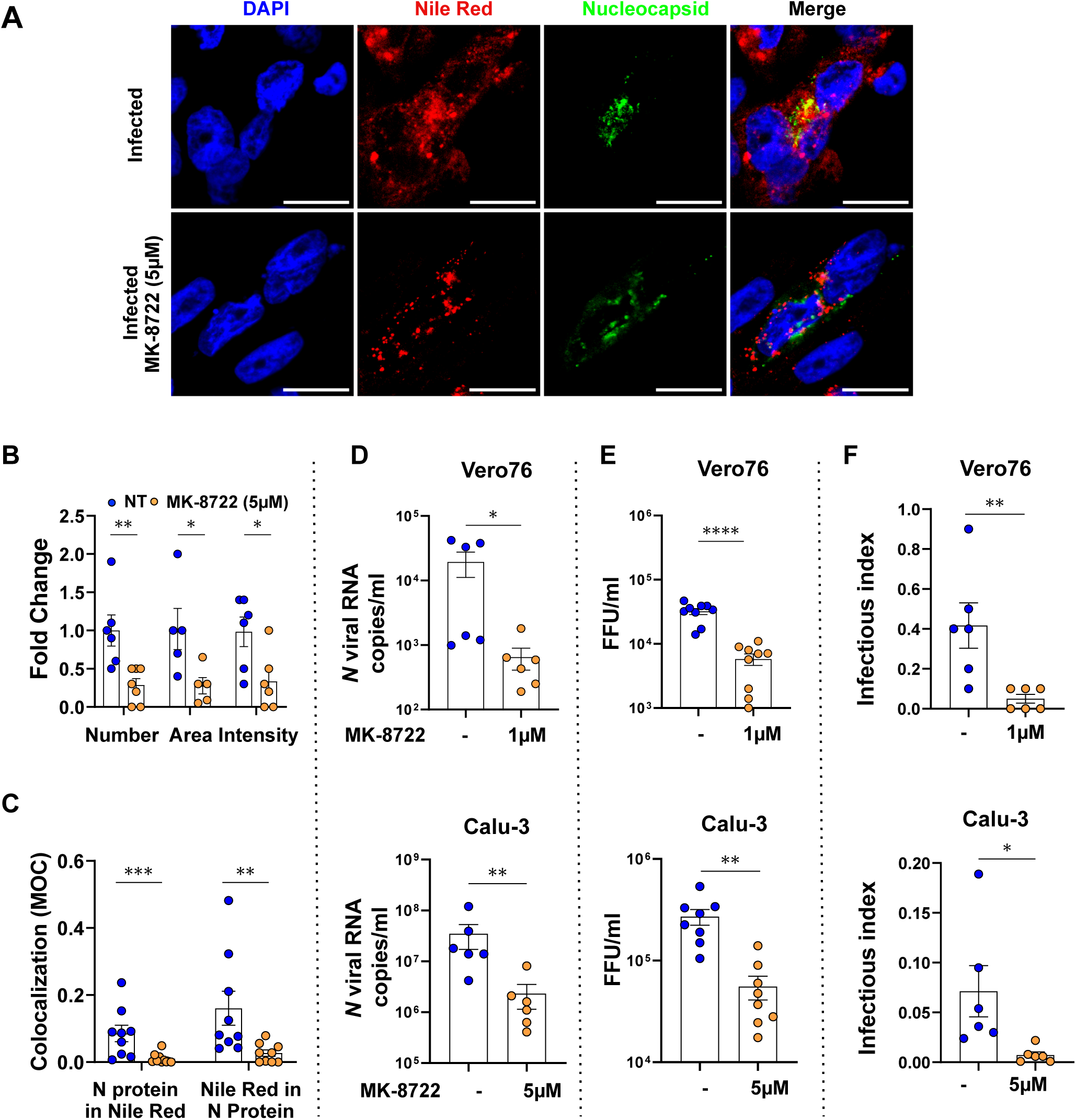
MK-8722 inhibition of SARS-CoV-2 is associated to decrease of lipid droplets and release of defective viral particles. **A: *Morphological analyses.*** Calu-3 cells were infected with the SARS-CoV-2 Alpha variant in the presence of a continuous treatment with MK-8722 (5 µM) or not. At 3dpi, lipids droplets, defined as indicated in the method section, were stained with Nile Red lipophilic dye before cells fixation. Nucleoprotein protein (Green) expression detected by immunofluorescence, Nile Red (Red) staining and nuclei staining with DAPI (Blue) were observed by confocal microscopy. One representative field out of 5 observed per condition are presented. Bar: 20 µm. Shown are images representative of n=3 independent experiments. **B: *Quantitative analysis of images*** shown in A using ImageJ of the number, area and intensity of lipid droplets (LD), as defined in the method section within infected cells, characterized by the expression of Nucleocapsid protein. Values were normalized to corresponding infected non-treated condition. n=3 independent experiments. **C: *Lipid droplets and N-protein colocalization*.** The fraction of lipid droplets colocalizing in Nucleocapsid staining and inversely were quantified using JACoP plugin from ImageJ software and corresponding Manders’ overlap coefficients (MOC) are presented. **D**: ***Viral genomes released***. Cells were infected with the Alpha SARS-CoV-2 variant in the absence or in the presence of a continuously treatment with MK-8722 at the indicated concentrations. Viral *N* RNA released by Vero76 cells at 1dpi (upper panel) and Calu-3 cells at 4dpi (lower panel) cells were quantified by RT-qPCR. n≥3 independent experiments. **E: *Viral titres released*** by Vero76 (upper panel) and Calu-3 (lower panel) cells treated as above in (**D**). n=3 independent experiments. **F: *Infectious index of released viruses*** determined as described in the methods section using matched viral titres and viral copy number values Vero76 (upper panel) and Calu-3 (lower panel) cells obtained as in (**D**) and (**E**). n=4 independent experiments. Student’s *t*–test and Mann-Whitney * *p*<0.05, ** *p*<0.01, *** *p*<0.001, **** *p*<0.0001.

SARS-CoV-2 is an enveloped virus; therefore lipid metabolism modification alters the lipid composition of the viral membrane, and in turn viral particles infectivity ^5,7–11^ We thus measured viral production by infected Vero76 and Calu-3 cells by quantification of secreted viral nucleocapsid *N* RNA by RT-qPCR. In line with the reduction of secreted viral *N* RNA in presence of the drug shown above (**Fig. 1E**), the level of viral *N* RNA secreted by both infected cell lines was reduced by 10-fold upon continuous MK-8722 treatment (**Fig. 3D**, 2±0.8×10^4^ vs 6±2×10^2^ and 3.4±1.7×10^7^ vs 2.3±1.2×10^6^ RNA copies/mL for Vero (upper) and Calu-3 cells (lower), respectively, *t*-test: *p*<0.05 for Vero76 and *p*<0.01 for Calu-3 cells). Accordingly, the amount of infectious viruses released by infected cells titrated in foci forming assay was statistically decreased by MK-8722 in Vero76 (from 3±0.3×10^4^ to 5.8±1×10^3^ FFU/mL) and Calu-3 cells (from 2.6±0.5×10^5^ to 5.5±1×10^4^ FFU/mL, *t*-test: *p*<0.05 for both cell types, **Fig. 3E**).

We next calculated an infectious index of released particles (described in the method section), which inversely correlates with virus infectivity. The drug reduced the frequency of newly produced infectious viruses by both cell lines (infectious index: 0.41±0.11 vs 0.05±0.02 in Vero76 cells (Mann-Whitney *p*<0.05), and 0.07±0.03 vs 0.007±0.003 in Calu-3 cells (Mann-Whitney *p*<0.05) in the presence and absence of the drug) (The drug reduced the frequency of newly produced infectious viruses by both cell lines **Fig. 3F**). This inhibitory effect of MK-8722 on the production of infectious particles could not be explained by a direct inhibitory role of MK-8722 present in the culture media on foci formation. Indeed, tested cell culture media (**Fig. 1B, S2B**) were highly diluted in the assay (at least 1,000 times) corresponding to a MK-8722 concentration < 0.005 µM, which is far under the limit of efficacy of MK-8722 in Vero76 cells. Furthermore, MK-8722 exposure was limited to the 2h inoculation. Our data suggests that the lipid metabolism alteration induced by MK-8722 counteracts SARS-CoV-2 replication and affects production and infectivity of released viral particles.

### MK-8722 enhances lysosomal degradation of viral components to promote antiviral immunity

As antiviral mechanism, MK-8722 could contribute to restoration of the autophagic flux (**Fig. 2**), which should result in targeting viral proteins to lysosomes. We first evaluated the lysosome distribution after MK-8722 treatment in infected cells. As a lysosomal marker, we used LAMP1 and characterized its distribution in infected Calu-3 cells upon MK-8722 treatment by immunolabelling and confocal microscopy in infected (**Fig. 4A**) and non-infected cells (**Fig. S5A**). SARS-CoV-2 infection significantly decreased LAMP1 relative expression in Calu-3 cells as expected (**Fig. 4A and S5A,** quantified in **Fig. S5B** as 0.47±0.06-fold change, ANOVA: *p*<0.05). This decrease likely reduced LAMP1^+^ lysosome size as infection inhibits autophagosome-lysosome fusion^22,24,25^. MK-8722 treatment during infection significantly restored LAMP1 relative expression to its level in non-infected non-treated condition (1.15±0.11-fold change, ANOVA: *p*<0.01 compared to infected non-treated condition; **Fig. S5B**).

**Figure 4:**
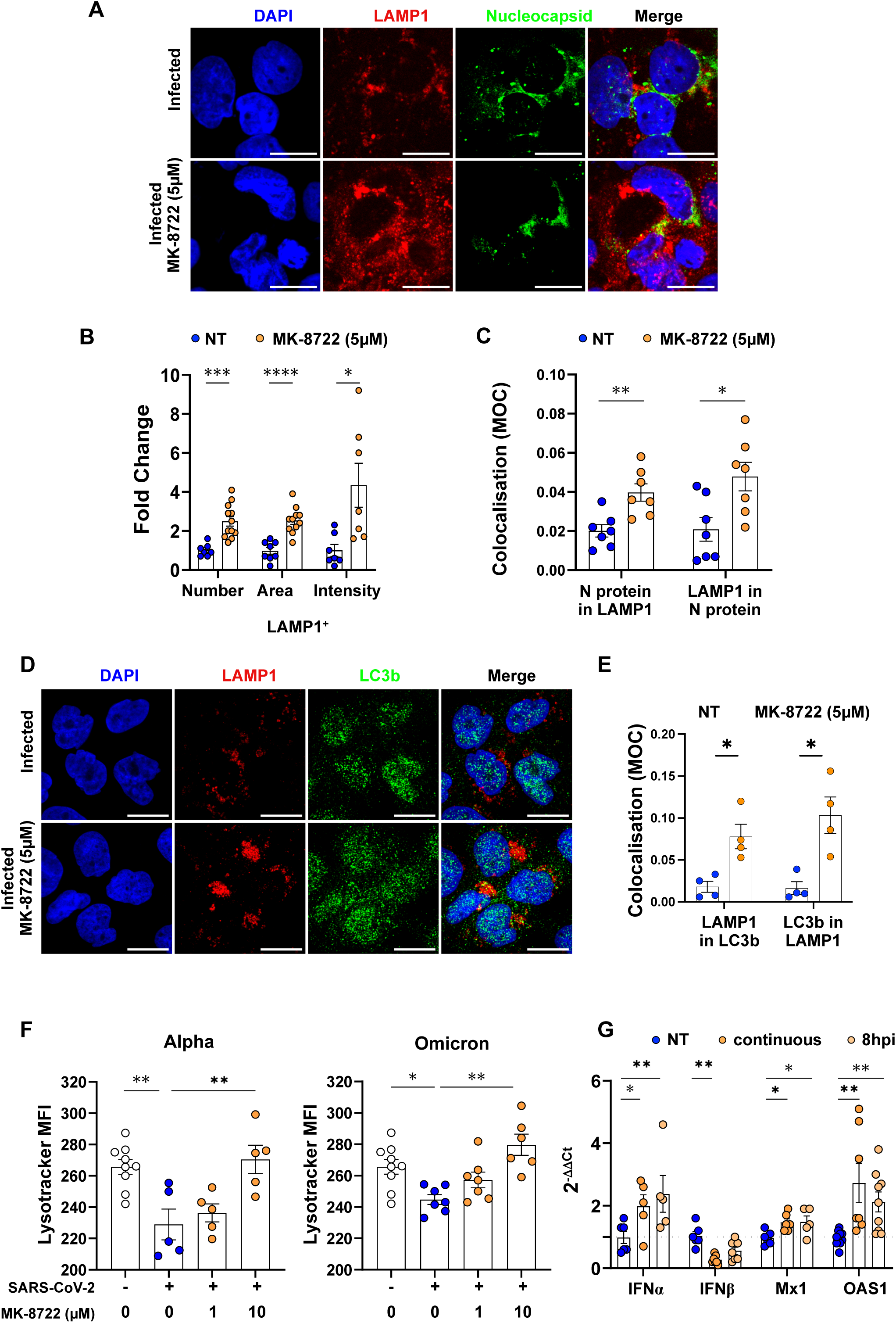
MK-8722 treatment increases the autophagic flux redirecting viral components to lysosomes and restores type I interferon responses.” **A: *Morphological analysis***: Calu-3 cells were infected with the SARS-CoV-2 Alpha variant in the presence of a continuously treatment with MK-8722 (5 µM) or not. At 3dpi, cells were then fixed and LAMP1 (Red) and Nucleocapside N (Green) protein expression was detected by immunofluorescence. Nuclei were stained with DAPI (Blue). Cells were observed by confocal microscopy. One representative field out of 5 observed per condition is presented. Bar: 20 µm. Images representative of n=3 independent experiments. **B: *Quantitative analysis*** of images show in A using ImageJ of number, area and intensity of lysosomes, identified as defined in the methods section, within infected cells, characterized by the expression of Nucleocapsid protein. Values were normalized to infected non-treated condition. n=3 independent experiments. **C: *LAMP1 and N protein colocalization*.** The fraction lysosomal objects colocalizing with Nucleocapsid staining and inversely were quantified using JACoP plugin from ImageJ software and corresponding Manders’ overlap coefficients (MOC) are presented. n=3 independent experiments. ***D: Morphological analysis***: Calu-3 cells were infected with the SARS-CoV-2 Alpha variant in the presence of a continuously treatment with MK-8722 (5 µM) or not. At 3dpi, cells were then fixed and LAMP1 (Red) and LC3b (Green) protein expression was detected by immunofluorescence. Nuclei were stained with DAPI (Blue). Cells were observed by confocal microscopy. One representative field out of 4 observed per condition is presented. Bar: 20 µm. Images representative of n=3 independent experiments. ***E: LAMP1 and LC3b protein colocalization*.** The fraction lysosomal objects colocalizing with LC3b foci staining and inversely, were quantified using JACoP plugin from ImageJ software and corresponding Manders’ overlap coefficients (MOC) are presented. n=3 independent experiments. ***F: Lysosomal activity.*** Calu-3 cells were infected with the indicated SARS-CoV-2 variant in the presence of indicated MK-8722 concentration added at 24hpi and cultured for another 24h. Lysosomal activity was then analysed using Lysotracker DND99 dye and quantified by flow cytometry. Shown are lysotracker MFI collected from n≥5000 cells analysed. n=3 experiment. ***G: IFN-I response***. Calu-3 cells were infected with SARS-CoV-2 Alpha variant in the presence of MK-8722 (5 µM) continuous treatment (dark orange) or initiated at 8hpi (light orange) or without treatment (Blue). IFN-I response was evaluated by quantifying *IFNα*, *INFβ*, *Mx1* and *OAS1* mRNA by RT-qPCR of indicated genes at 1dpi. Data represent the expression of each mRNA relative to *β-actin* mRNA levels, and expressed following the 2^-ΔΔCt^ method. n=3 independent experiments. ANOVA and Mann-Whitney * *p*<0.05, ** *p*<0.01, *** *p*<0.001, **** *p*<0.0001.

We then characterized LAMP1^+^ lysosome parameters, as defined in the method section. MK-8722 significantly increased LAMP1^+^ lysosome number, area and specific intensity, indicative of their enhanced activity in infected cells, which were identified by the expression of viral N protein (**Fig. 4B**, 2.5±0.25 and 2.53±0.2 and 4.3±1.1-fold change over infected non-treated cells, *t*-test: *p<*0.001, *p<*0.05 and *p<*0.0001, respectively). When colocalization of viral N protein and LAMP1 was quantified in infected cells, MK-8722 treatment resulted in higher MOC as compared to infected non-treated cells (N protein in LAMP1 signal: 0.02±0.003 vs 0.05±0.004, Mann-Whitney *p<*0.01; and LAMP1 in N protein signal: 0.021±0.006 vs 0.048±0.007, Mann-Whitney *p<*0.05, **Fig. 4C**). To assess directly the impact of MK-8722 on the autophagic flux, Calu3 cells were infected by SARS-CoV-2 without and with MK-8722 (5uM), double labelled with LC3b and LAMP1, and co-localisation of the two markers quantified (**Fig.4D, E**). MK-8722 treatment, compared with no treatment, increased LC3b colocalization in the LAMP1 compartment as shown by the increase in MOCs in treated versus non treated infected cells (LAMP1 signal in LC3b signal 0.078±0.014 vs 0.01±0.006, Mann-Whitney p<0.05, and LC3b in LAMP1 signal 0.04±0.02 vs 0.015±0.007 Mann-Whitney p<0.05). These results indicate that MK-8722 restored the autophagic flux that had been interrupted by SARS-CoV-2 infection in Calu3 cells.

Lysosomal activity was quantified after Lysotracker staining by flow cytometry in Calu-3 cells infected for 24h and treated with MK-8722 for an additional day. MK-8722 increased lysosomal activity similarly after infection by both Alpha and Omicron variants (Lysotracker MFI: 286±12 and 289±9.6 in MK-8722 treated vs 253±34 and 251±6 in non-treated cells, respectively for each variant, ANOVA: p<0.05 at least, **Fig. 4F**). This result indicates that MK-8722 restores the autophagic flux to address viral components to the lysosome, where they are degraded.

### MK-8722 restores the INF I pathway

The lack of type-I IFN response is a major immune parameter contributing to COVID19 disease ^29,41,42^. Completion of autophagy will most likely result in the degradation of viral proteins, which escape RIG-I sensing thereby restoring IFNα and IFNβ production and associated downstream ISG such as OAS1 and Mx1. We thus investigated IFN-I response in infected Calu-3 cells after continuous or post-infection treatment with MK-8722 (**Fig. 4G**). Compared to untreated condition, continuous and post-infection treatment with MK-8722 increased *IFNα* mRNA expression by 100±38% and 140±58% (Kruskal-Wallis: *p<*0.05 for both treatments), while that of *IFNβ* decreased by 73±5% and 45±11% for continuous and post-infection treatment, respectively (Kruskall-Wallis: *p*<0.01 and ns respectively). Regarding *ISG* mRNA expression, *Mx1* level increased by 1.46±0.12-fold and by 1.49±0.19-fold for continuous and post-infection treatment, respectively (Kruskal-Wallis: <0.05 both, respectively), while *OAS1* mRNA expression increased by more than 2-fold in all treated conditions (2.8±0.6-fold and 2.1±0.3-fold, respectively, Kruskal-Wallis: *p*<0.01). Altogether, MK-8722 treatment, even if initiated post-exposure, can drive early IFN-I expression and downstream ISG, which, by controlling infection, could limit severe outcome in patients ^26,27,29,41,43^.

### MK-8722 *in vitro* treatment does not alter T cell response against SARS-CoV-2 in vaccinated individuals

T cell response is crucial to control SARS-CoV-2 infection ^44–46^, and relies on metabolism reprogramming of the effector cell. We therefore investigated the impact of MK-8722-mediated AMPK activation on the T cell response using primary cells obtained from peripheral blood of healthy individuals. We found that MK-8722 alone compared to DMSO at identical concentration did not activate T-cell from healthy donors (**Fig. S6A-B**). We then investigated if MK-8722 exposure affected T cell responses upon TCR engagement. Healthy donor PBMCs were stimulated with anti-CD3 and anti-CD28 (CD3/CD28, TransAct) to mimicking TCR engagement, and treated with MK-8722 for 6h. Intracellular IFNγ^+^ and CD107a^+^ frequencies were then analysed in CD4^+^ and CD8^+^ T cells. As expected, CD3/CD28 stimulation increased both IFNγ^+^ and CD107a^+^ frequencies in both CD4^+^ and CD8^+^ T cells (**Fig. 5A**, 5.6±1.5% and 7.5±1.9% vs 0.6±0.2% and 1.1±0.2% for CD4^+^ T cells, and 4.9±1.5% and 2±0.3% vs 0.68±0.1% and 0.3±0.06% for CD8^+^ T cells, for IFNγ^+^ and CD107a^+^ frequencies respectively, *t*-test: *p<*0.05 at least). The concentrations of MK-8722 tested had no negative impact on the expression of activation markers, and even showed a trend towards increased activation with MK-8722 treatment. These results indicate that MK-8722 does not affect degranulation and IFNγ responses and is thus compatible with the establishment of a T cell-dependent protective immune response.

**Figure 5:**
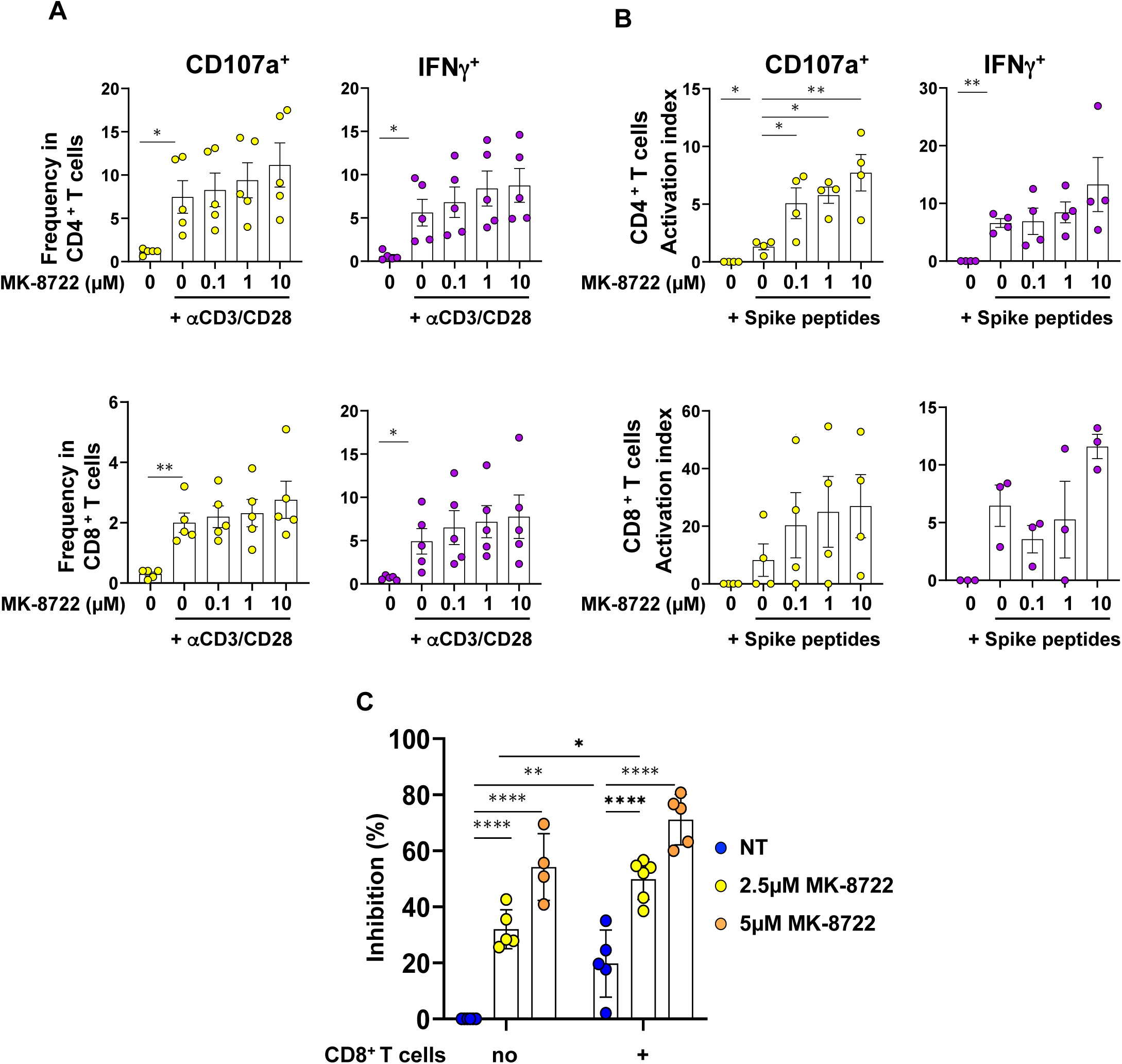
MK-8722 treatment does not alter T cell response from vaccinated individuals against SARS-CoV-2 *in vitro*. **A:** PBMCs from healthy SARS-CoV-2 vaccinated donors depleted in CD14 cells were stimulated for 6h with anti-CD3 and anti-CD28 antibodies in the absence or in presence of MK-8722 (5 µM). IFNγ^+^ and CD107a^+^ cell frequencies within CD4^+^ and CD8^+^ T cell populations were evaluated by flow cytometry. n=5 independent donors analysed in independent experiments. **B:** CD14-depleted PBMCs from corresponding donors in **(A)** were stimulated for 6h with a T cell SARS-CoV-2 spike peptide pool (Peptivator, Miltenyi), IFNγ^+^ and CD107a^+^ cell frequencies were quantified as in (A). Activation levels are presented as activation index calculated as described in the methods section. n≥3 independent donors analysed in independent experiments. **C:** Caco2 cells were infected 2h with the Alpha variant of SARS-CoV-2 and treated post exposure (8hpi) _+_ with MK-8722 (5 µM) alone or together with CD8 T cells from SARS-CoV-2 vaccinated individuals. Infection was evaluated by quantification of Spike expression in cells at 2dpi by flow cytometry and presented as inhibition of Spike expression frequency. n=5 independent donors performed in independent experiments. Data presented are mean ± SEM. ANOVA and *t*-test: * *p*<0.05, ** *p*<0.01, *** *p*<0.001, **** *p*<0.0001

We next tested whether MK-8722 altered antigen-specific T cell stimulation. As a surrogate of SARS-CoV-2 antigen after infection, we used peptides covering the immunodominant sequence domains of the SARS-CoV-2 Spike (Peptivator peptide pool (Miltenyi)) to stimulate CD14-depleted PBMCs and measure IFNγ^+^ and CD107a^+^ frequencies in both CD4^+^ and CD8^+^ primary T cells. We defined an activation index as the level of activation induced by the peptide pool relative to the corresponding Transact level. For these experiments, cells were collected from healthy individuals after summer 2021 when most of them had been immunized at least two times with spike-based SARS-Cov-2 vaccine and consequently mounted a Spike-specific T cell response. Accordingly, Spike peptide-pool stimulated only CD4^+^ T cell response although in a limited manner (**Fig. 5B**, activation index: 6.6±0.8 for IFNγ and 1.4±0.3 for CD107a, *t-test: p*<0.01 and *p*<0.05). As for CD3/CD28 stimulation, MK-8722 stimulation showed a trend towards increase all the tested T cell responses although only CD107a activation index for CD4+ T-cells was statistically higher after MK-8722 treatment (i.e. 6.6±0.8 vs 13.3±4.7 for 10 µM MK-8722, ANOVA *p*<0.01). Stimulation of CD14-depleted PBMCs with a SARS-CoV-2 Nucleocapsid peptide pool gave similar results, even though the peptide pool alone failed to activate a T cell response (**Fig. S6C**). These data suggests that if used as a drug, MK-8722 will likely not prevent the establishment of a T cell-specific response against SARS-CoV-2 *in vivo*.

We finally evaluated whether MK-8722 and SARS-CoV-2-specific CD8^+^ T cells could cooperate to block infection of mucosal epithelial cells. However, Calu-3 cells being HLA-A2 negative, they could not be use to present viral antigen to HLA-A2+ CD8+ T cells. For this reason instead of Calu-3 cells, we used the epithelial Caco2 cells competent for IFN responses, which expressed HLA-A2 (not shown) and are susceptible to SARS-CoV-2 infection Therefore, CD8^+^ T cells were obtained from HLA-A2^+^ healthy donors obtained after the summer 2021. Post-exposure treatment with MK-8722 inhibited infection of Caco2 cells in a dose dependent manner, as shown in **Fig. 5C** (by 32±3% and 54±6%, at 2.5 and 5 µM respectively (ANOVA: *p*<0.0001 for both conditions, see **Fig. S7** for the gating strategy). In the presence of CD8^+^ T cells, infection of Caco2 cells was reduced by 19.8±5% (ANOVA: *p*<0.01). Moreover, the combination of CD8^+^ T cells with MK-8722 increased the inhibition of infection in a dose-dependent manner (50±3% and 71±4% inhibition in the presence of CD8+ T cells and 2.5 or 5 µM MK-8722, respectively) compared to the inhibition of infection in the sole presence of CD8+ T cells (19.8±5% ANOVA: *p*<0.0001 both, respectively) and in the presence of MK-8722 only (32±3% and 54±6% in the presence of 2.5 or 5 µM MK-8722, *t*-test *p*<0.05 and ns for 2.5 and 5 µM, respectively) although no synergy was observed. Altogether, our results confirm that MK-8722 treatment does not impair the SARS-CoV-2-specific CD8^+^ T cell-mediated response against mucosal Caco2 cell infection, but rather that both act in combination to reduce cell infection.

## DISCUSSION

To date, antivirals against SARS-CoV-2 infection without significant side effects have yet to be characterised. Here, we provide evidence that the direct pan-AMPK allosteric activator MK-8722 is a strong antiviral drug candidate against SARS-CoV-2 infection, as treatment with MK-8722 in the μM range reduces infection of lung cells without altering the cellular immune response.

Previous studies have suggested that AMPK activation could inhibit viral infections ^34^, as viral infection prevents AMPK activity. In the case of SARS-CoV-2 infection, AMPK activity is reduced and furthermore, autophagy is blocked ^18^, while inhibition of either AMPK by Compound C or ULK1 by MRT68921 increases viral particle release ^18,47^. We show here that the blockade of AMPK activation upon infection can be reversed by MK-8722, the pharmacological allosteric pan-activator of AMPK, which blocks infection at a µM concentration, in agreement with the predicted role of AMPK activity on SARS-CoV-2 infection^48^ and with MK-8722 action on infection by other viruses^49^.In line, metformin, a drug approved by FDA since 1994, and the adenosine analogue AICAR that activates AMPK indirectly can inhibit replication of SARS-CoV-2 as well as Flaviviruses, but at a concentrations of 10 and 1 mM, respectively ^35,47^. These AMPK-sensitive viruses replicate all in viral factories, disturbing lipid synthesis and escaping autophagy ^50,51^. AMPK activation by metformin at high concentration (10mM) inhibit SARS-CoV-2 replication in vitro ^47^. However, AMPK activation with 5-amino-imidazolecarboxamide riboside (AICAR), a non-metabolised analogue of AMP able to activate AMPK, used at 25 μM is unable to inhibit SARS-CoV-2 infection ^18^ while at 1 mM has been proven to reduce by 10-fold viral ^47^ suggesting that AMPK needs to reach an activation threshold to inhibit viral replication. Conversely, NUAK2, an AMPK-related kinase, was reported to stimulate viral replication in A549 and Calu3 cells ^52^. Altogether our results, in agreement with the literature, indicate that AMPK-dependent antiviral activity is restricted to AMPK-members only, confirming their distance with AMPK-related kinases such as NUAK2 ^53^. Despite observational data suggesting an association between metformin use and prevention of severe COVID-19 outcomes, metformin did not significantly lower risk of hospitalization and mortality due to SARS-CoV2 infection in recent randomized controlled trials ^54^. Furthermore, in vivo, metformin mainly targets the liver and the gastrointestinal tract ^55^ and therefore could have limited effect in the upper airway, the SARS-CoV-2 infection site. In contrast, MK-8722, as systemic drug, may reach these tissues with minimal side effects, as a daily treatment in diabetic Non-human primates (NHPs) with MK-8722 (10 mg/kg) for a month induced only a limited and reversible cardiac hypertrophy ^36^. The MK-8722 antiviral treatment we report is efficient *in vitro* not only when applied from the onset of infection but also after infection. Therefore, MK-8722 might be tested as post-exposure prophylaxis against SARS-CoV-2, thereby limiting potential severe outcomes.

Mechanistically, we demonstrated here that AMPK activation by MK-8722 reduces lipid synthesis through the inhibition of ACC by acting on the limiting step of fatty acid production and, consequently, of phospholipid synthesis. This inhibition leads to a reduction in the formation of lipid droplets and a significant reduction in the infectivity of viral particles. Lipid synthesis is required to extend endoplasmic reticulum for the genesis of the viral factories and to generate the viral membrane of SARS-CoV-2, which is an enveloped virus. Changes in lipid content of viral particles have been shown to impact infectivity ^5,7,9,11^. AMPK-mediated inhibition of ACC by MK-8722 treatment likely modifies all membrane lipid species in SARS-CoV-2 infected cells ^56^, in turn affecting viral membrane lipid composition and likely the lower virus infectivity we observed. The modification of the cellular lipid content by MK-8722 treatment might also block the release of pro-inflammatory molecules otherwise induced by infection, such as IL-6 and LTB4 ^12,57^. SARS-CoV-2 affects also cholesterol metabolism ^7,58,59^ and patients under statins who contracted SARS-CoV-2 developed better T cell responses and showed lower grade of pulmonary symptoms ^60,61^. AMPK activation also inhibits HMG-CoA-reductase, the major enzyme for cholesterol synthesis. MK-8722 treatment might therefore be beneficial for COVID-19 patients through multiple mechanisms ^7,14,17^. The molecular mechanisms underlying the changes in the lipid spectrum and cholesterol content in infected cells following treatment with MK-8722 remain to be understood.

Increased activation of ULK1 and expression of Sequestosome-1/p62 upon MK-8722 treatment *in vitro* reflects a restoration of the autophagy flux and the targeting of viruses to lysosome for degradation. This mechanism is otherwise blocked by viral replication in an ORF3- and ORF7a-mediated process, which inhibits HOPS complex-mediated assembly of the SNARE complex required for autolysosome formation ^22,24,25^. Sequestrosome-1/p62 is an autophagy cargo receptor that plays a key role in mediating the formation of autophagosomes and autophagic clearance of intracellular protein aggregates by its binding to LC3b. A modification of p62 expression and LC3b binding capacity upon SARS-CoV-2 infection has been observed *in vitro* ^20,22,25^. Together, these data suggest that SARS-CoV-2-mediated disruption of p62 expression and/or function impairs fusion of phagosome with lysosome ^20,22,62,63^. The increased p62 expression we observed upon MK-8722 treatment together with the increased ULK1 activation will likely result in p62 activation, in turn promoting phagolysosomes formation as we reported. Convergence of Beclin-Atg14 together with P62 activation could stimulated the selective clearance of viral components, in a process called virophagy ^64–66^. We thus propose that AMPK pharmacological activation induces virophagy and is responsible, at least partially, for its antiviral effect. Furthermore P62 also induces proteasomal degradation^67^, and its increased expression could contribute in the antiviral activity of MK-8722.

The restoration of autophagy by MK-8722 we report, and thereby degradation of viral components, will improve MHC-I presentation and the cellular cytotoxic response ^68^, contributing to the therapeutic value of MK-8722 treatment. Furthermore, defective viral particles, which are produced upon MK-8722 treatment, might be detected by innate immune sensors such as MAVS ^69^, promoting a type-I IFN response and protection from SARS-CoV-2 infection ^28^. MK-8722 treatment of infected cells also unlocks the IFN-I response, as shown by the increase in *IFNα* and both ISG *Mx1* and *OAS1* mRNA we observed. This IFN-I response most likely associates with a better control of infection and lowers pathology score ^26,27,29,41,70^. The induction of IFNβ by poly I:C is only transient, with its mRNA returning even to a level below the basal one ^71^. Here, continuous but not post infection treatment with MK-8722 may similarly result in a transient accumulation of *IFNβ* mRNA, the final level of which is statistically lower than that observed in the absence of treatment. This faster IFN-I response could enhance protection of individuals against SARS-CoV-2, since a delay in IFN-I response is associated with disease severity ^29,41^ and post infection treatment of severe COVID with IFNα2b significantly improved patient outcome ^42^.

Finally, MK-8722 could also enhance the response of SARS-CoV-2-specific CD8^+^ T cells at the site of infection which is crucial for protection in NHP and in humans ^44–46^. Optimal CD8^+^ T cells responses at mucosal level can be boosted by IFN-I, which showed vaccine adjuvant-like properties in strategies against other infectious diseases and cancer ^72–75^. MK-8722 treatment, which stimulates the IFN-I pathway upon infection, as shown here, could therefore contribute to improve the CD8^+^ T cells response. Accordingly, *in vitro*, MK-8722 does not antagonise but rather slightly enhance the T cell response and when combined with CD8^+^ T cells, resulting treatment increases inhibition of infection compared to either treatment alone. Further studies on the crosstalk between CD8^+^ T cells, phagocyte and infected epithelial cells will clarify how MK-8722 modulates the CD8^+^ T cell response. The antiviral effect of AMPK activation in primary lung reconstruction and preclinical models such as hamster remains to be tested.

In summary, to optimize safely antiviral treatment against SARS-CoV-2, as summarized in the graphical abstract, we propose here to target the energy sensor AMPK using small molecule pan-AMPK allosteric activators (such as MK-8722), rather than targeting the virus itself. The antiviral mechanism of MK-8722 relies on the reversion of three major pathways affected by SARS-CoV-2 infection, namely lipid metabolism, autophagy and IFN-I response. Therefore, to escape MK-8722 and generate resistant and more pathogenic variants, SARS-CoV-2 will need to acquire a higher number of resistance mutations than a drug targeting on a single pathway. Furthermore, by increasing the SARS-CoV-2 CD8^+^ T cell response, MK-8722 treatment would result in a faster response against infection and better outcome. *Coronaviruses* but also *Flaviviruses* replicate within viral factories and/or disturb lipid metabolism, hijack autophagy and IFN-I response. We therefore propose that MK-8722 could act as antiviral against other *Coronaviruses* and *Flaviviruses*, ensuring a larger available therapeutic arsenal against future emerging virosis.

## Supplementary Figure Legends

**Supplementary Figure S1:**
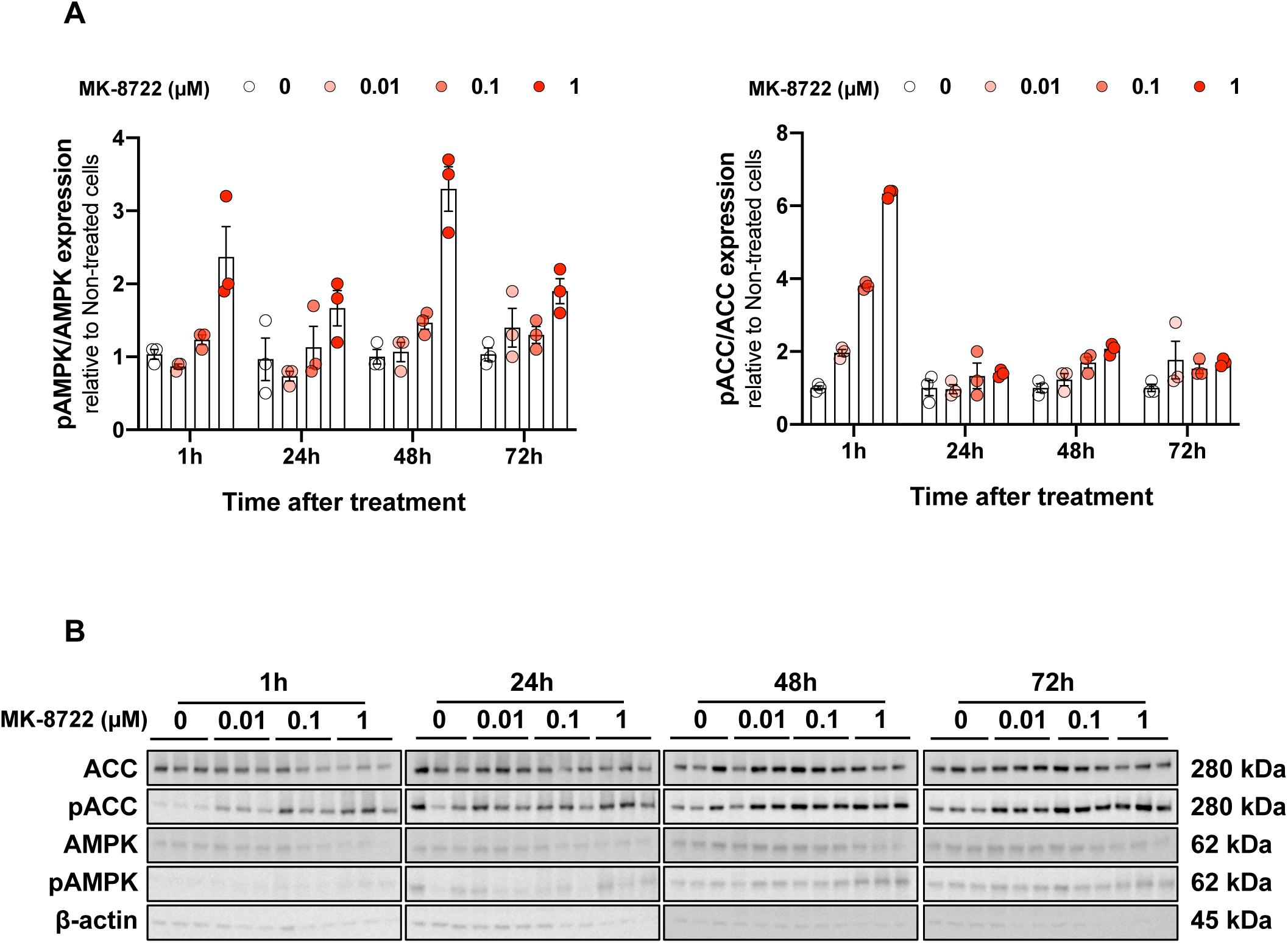
Kinetic of AMPK activation and downstream ACC phosphorylation. evaluated by western-blot after MK-8722 stimulation of Vero76.

**Supplementary Figure S2:**
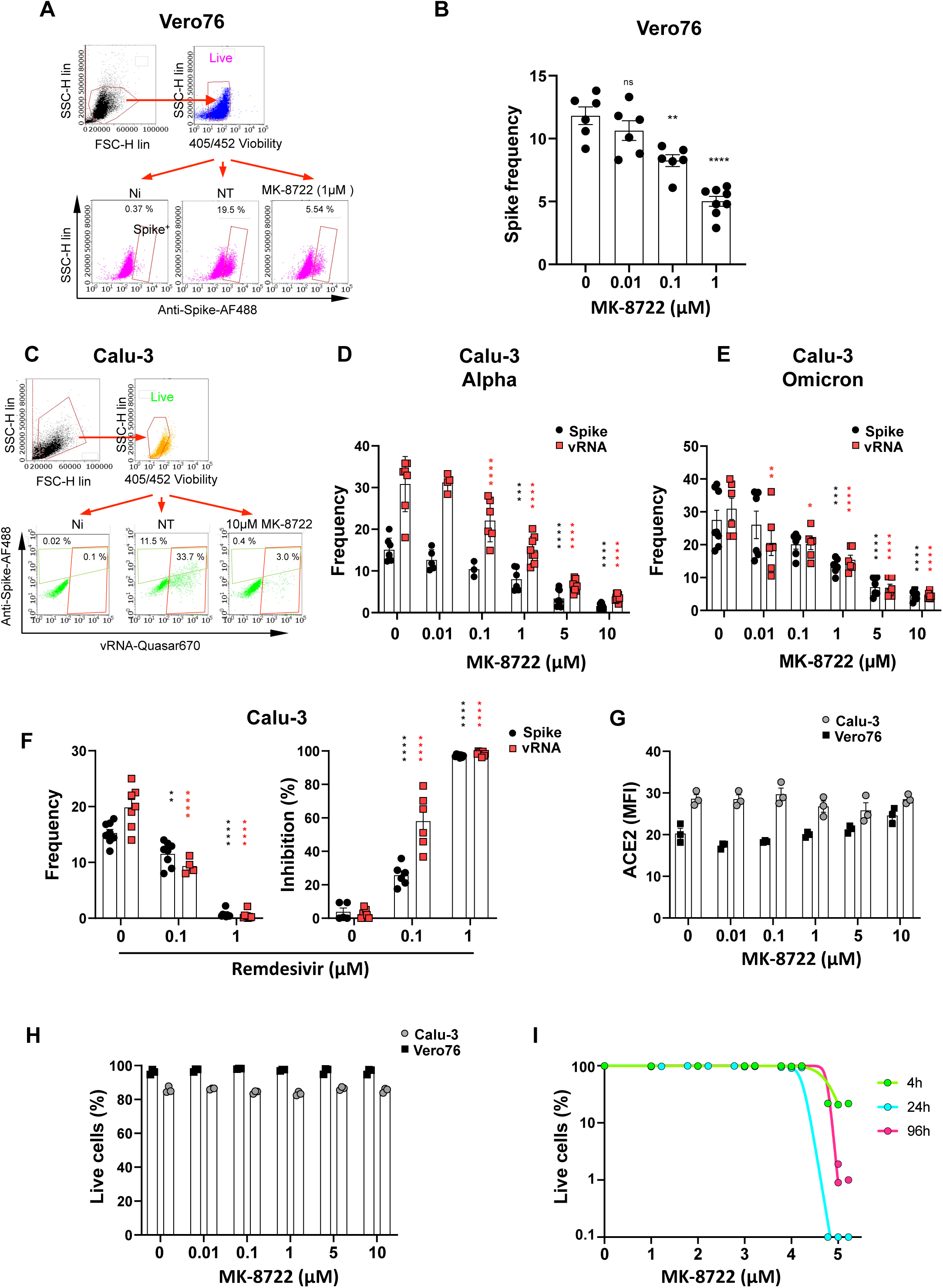
MK-8722 inhibits SARS-CoV-2 infection. **A:** Gating strategies used to analyse infection inhibition by MK-7822 treatment in Vero76 cells in Figure 1B. **B:** Cells were treated as indicated in Figure 1B. Frequencies of infection in corresponding experimental sample shown in Figure 1B were measured by SARS-CoV-2 Spike detection in Vero76 cells. Shown are mean ± SEM. n=4 independent experiments. **C:** Gating strategies used to analyse infection inhibition by MK-7822 treatment in Calu-3 cells in Figure 1C. **D-E:** Cells were treated as indicated in Figure 1B and C. Frequencies of infection in corresponding experimental sample shown in Figure 1B and C were measured by dual detection of Spike (Black) and viral RNA (Red) using Fish-flow in Calu-3 cells infected by Alpha **(D)** or Omicron **(E)** variant, quantified by flow cytometry. Shown are mean ± SEM. n=4 independent experiments. **F**: Inhibition of infection by Remdesevir. Calu-3 cells were treated with Remdesivir (0.1 µM or 1 µM) 1h prior and during the 2h of inoculation with SARS-CoV-2 Alpha variant. After virus removal, treatment with Remdesivir at indicated concentration was continued. At 4dpi, infection was quantified by dual detection of Spike (Black) and viral RNA (Red) in Fish-flow quantified by flow cytometry. Shown are mean ± SEM of Spike and vRNA frequency (left) and infection inhibition (right). n=3 independent experiments. ***G-H*:** ACE2 expression and viability were evaluated in Vero76 cells after 24h or in Calu-3 cells after 4days of MK-8722 continuous treatment (1 µM or 5 µM respectively) by flow cytometry. ACE2 expression is expressed as MFI (***G***) and viability as frequency of cells non-stained by the amine-reactive dye Viobility (***H***). n=3 independent experiments. ***I:*** Evaluation of MK-8722 over time and concentration in Calu-3 cells. n=3 independent experiments. Shown are mean ± SEM. ANOVA: * *p*<0.05, ** *p*<0.01, *** *p*<0.001, **** *p*<0.0001.

**Supplementary Figure S3:**
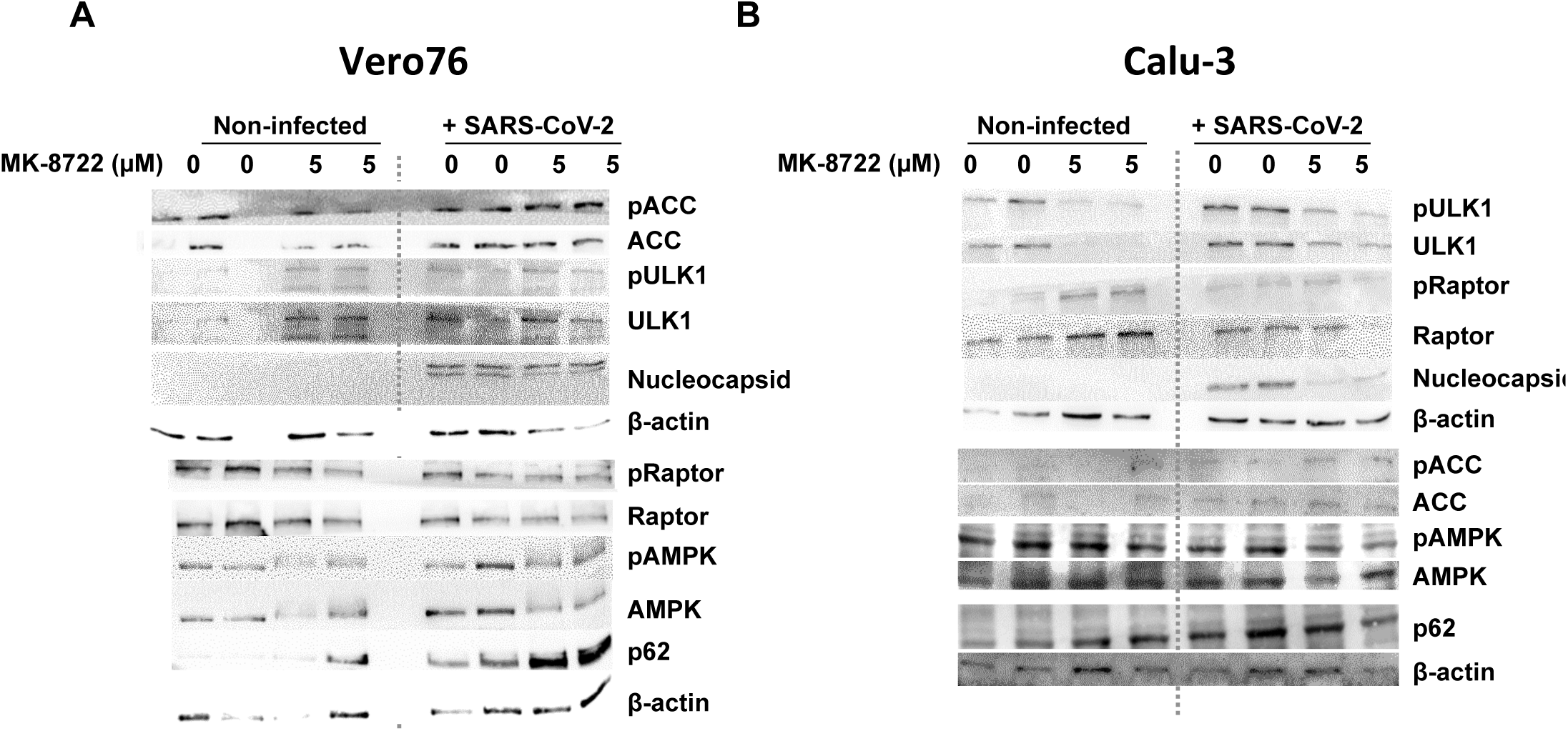
Evaluation of MK-8722 activity on cellular pathways downstream AMPK activation by western blot of ACC, ULK1, Raptor and AMPK phosphorylation, and Nucleocapsid protein N and autophagy cargo receptor p62 expression in Vero76 **(A)** and Calu-3 cells **(B)** infected or not and continuously treated or not with MK-8722 (5 µM) as indicated. The image shown is a montage of non-adjacent from same western blots experiments used to evaluate each protein and, when indicated, corresponding phosphorylated form.

**Supplementary Figure S4:**
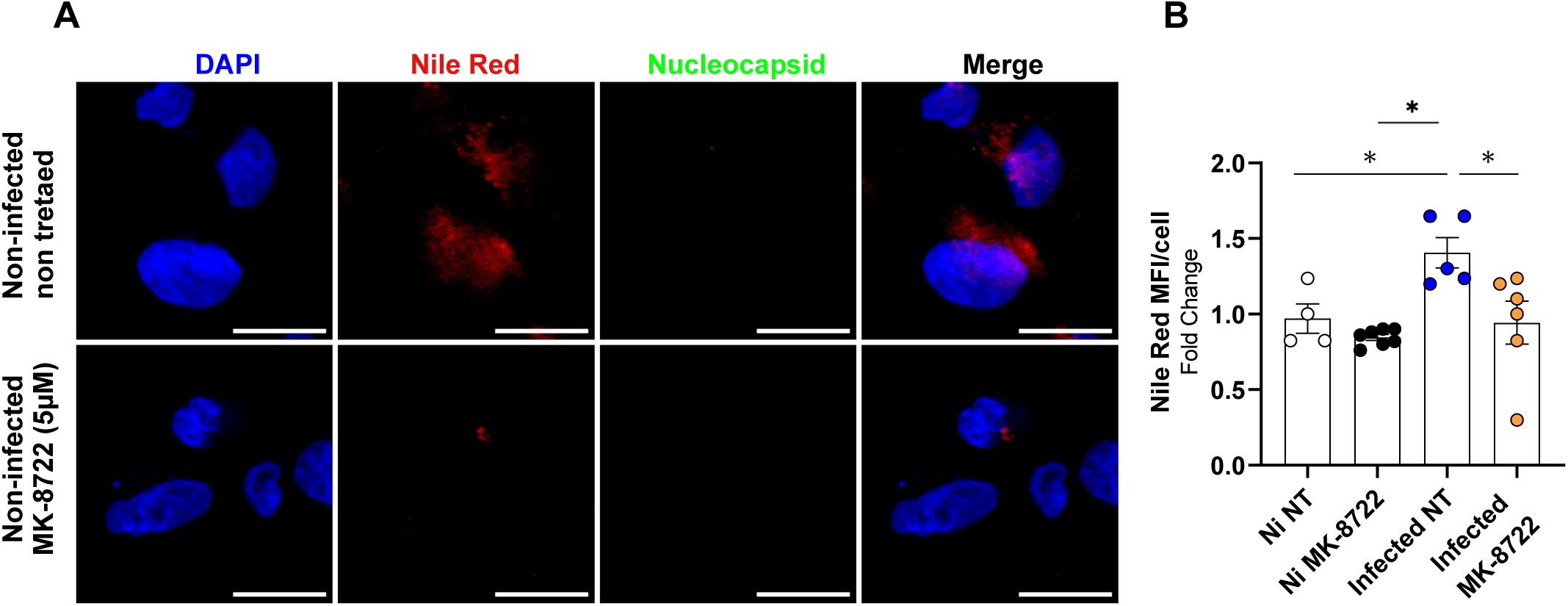
MK-8722 treatment decreases lipid droplet distribution in uninfected Calu-3 cells. **A:** Cells were left uninfected and treated with MK-8722 (5 µM) or not and analyzed as indicated in Figure. Representative fields for Nile Red (Red) and Nucleocapsid N protein immunodetection (Green) imaged by confocal microscopy are shown. Nuclei were stained with DAPI (Blue). Bars: 20 µm **B:** Cells were treated and labelled as described in Figure 3. Total intensity of total Nile Red staining per cell was calculated in ImageJ for non-infected non-treated (Ni-NT), infected non-treated (Infected-NT) and Infected MK-8722 treated conditions. Values were normalized to the mean of non-infected non-treated condition (Ni-NT). Shown are mean ± SEM. n= 3 independent experiments. ANOVA: *p<*0.05.

**Supplementary Figure S5:**
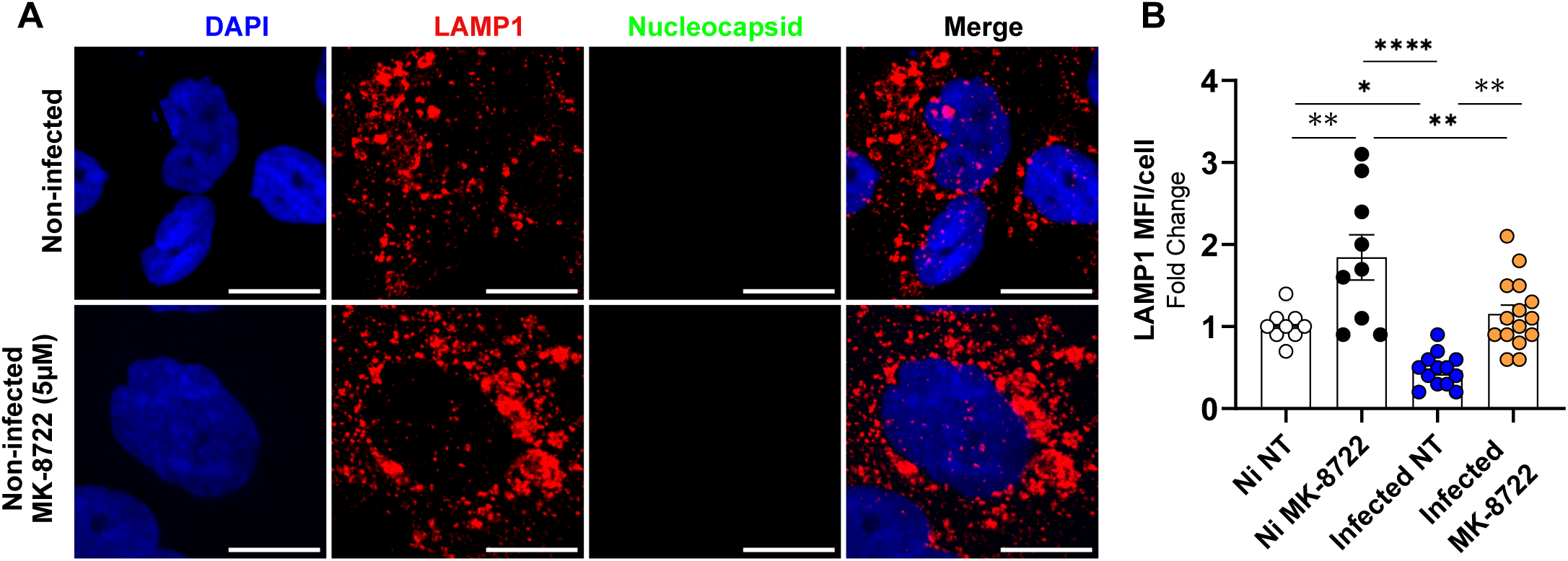
MK-8722 impacts lysosomal parameters in Calu-3 cells. **A:** Cells were left uninfected but treated with MK-8722 (5 µM) or not and analysed as indicated in Figure 4. Representative fields for LAMP1 (Red) and Nucleocapsid N protein (Green) immunodetection imaged by confocal microscopy are shown. Nuclei were stained with DAPI (Blue). Bars: 20 µm **B:** Cells were treated and labelled as described in Figure 4. Total intensity of total LAMP1 staining per cell was calculated in ImageJ for non-infected non-treated condition (Ni-NT), infected non-treated (Infected-NT) and treated with 5 µM MK-8722 (Infected-MK-8722). Values were normalized to corresponding non-infected non-treated condition. n≥3 independent experiments. Shown are mean ± SEM. ANOVA: ** *p<*0.01 and *** *p<*0.001.

**Supplementary Figure S6:**
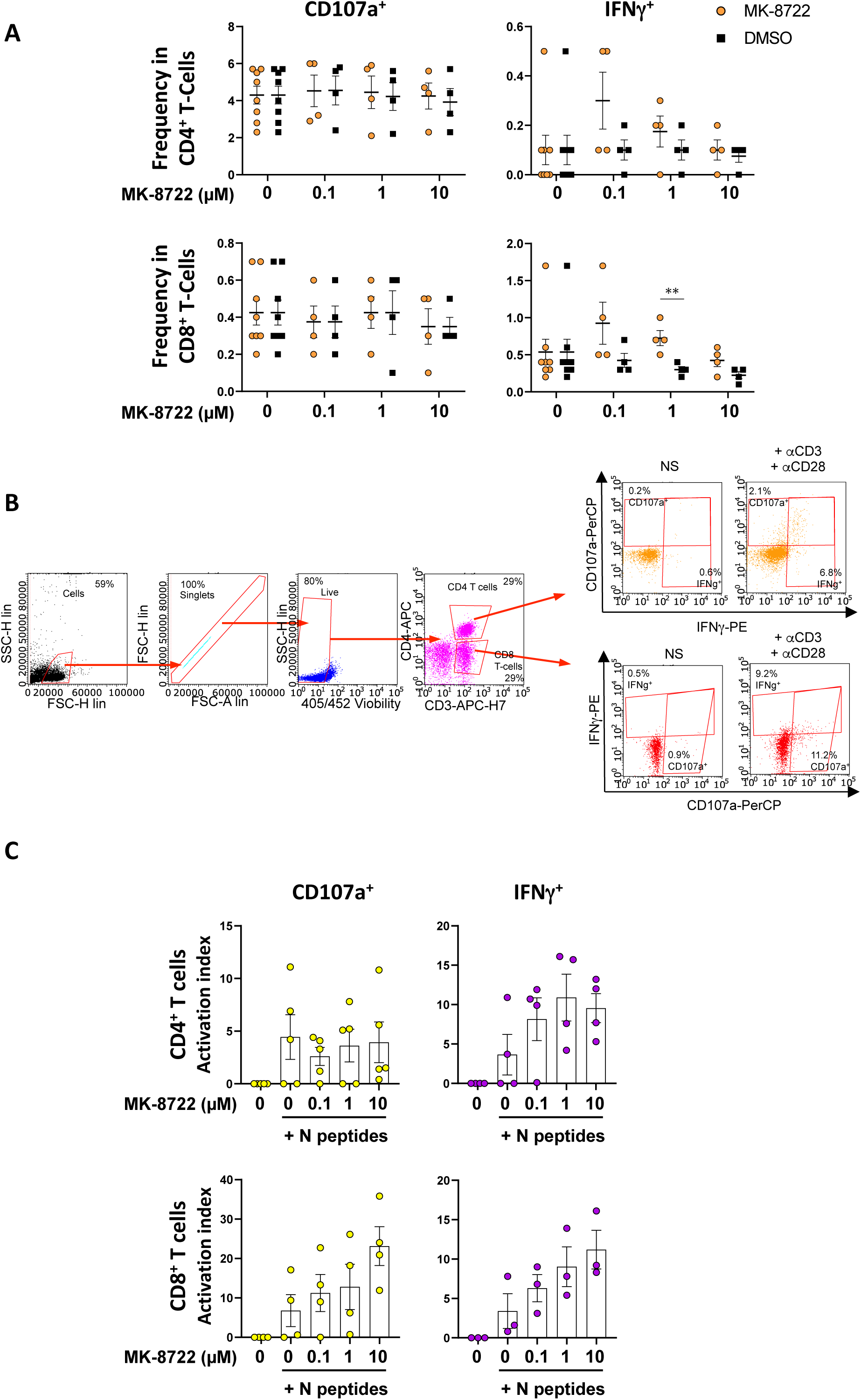
Evaluation of nucleocapsid-specific T cell response in SARS-CoV-2 vaccinated individuals upon MK-8722 treatment ex vivo. **A:** PBMCs from healthy donors were cultivated for 6h indicated concentration of MK-8722 or corresponding DMSO serving as carrier. CD4 and CD8 T cells activation (CD107a and IFNg expression) was monitored by flow cytometry. **B:** Gating strategy to investigate CD4^+^ and CD8^+^ T cell responses stimulation after activation with anti-CD3/anti-CD28 antibodies (Transact) with or without MK-8722 treatment as analysed in Figure 5A. **C:** CD14-depleted PBMCs corresponding to donors used in Figure 5 were stimulated as described in Figure 5B but with T cell Nucleocapsid peptides (Peptivator, Miltenyi), and monitored by flow cytometry for IFNγ and CD107a expression. Activation levels are presented as activation index as described in the methods section. n≥3 independent donors.

**Supplementary Figure S7:**
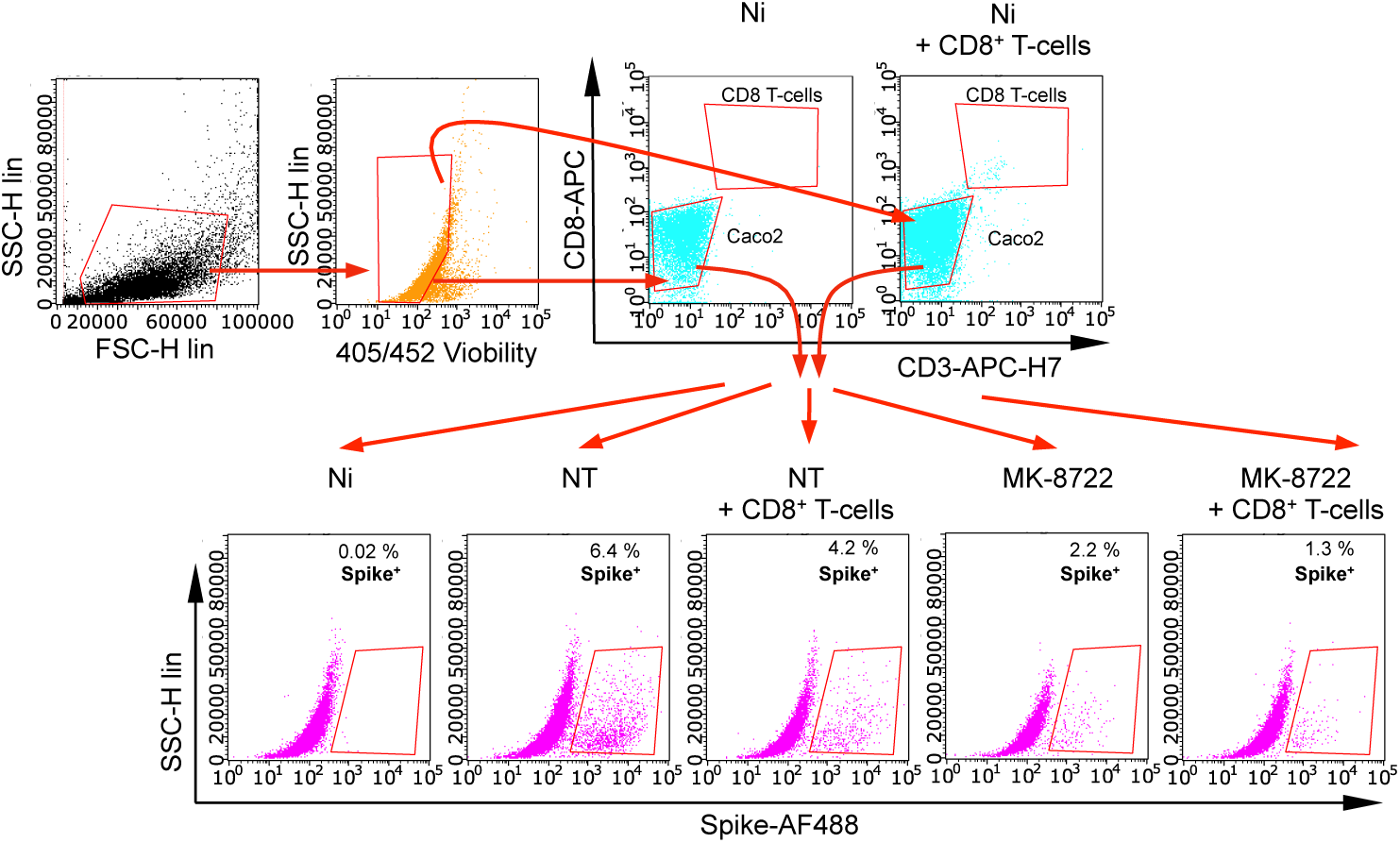
Gating strategy for combined MK-8722 treatment and CD8+ T cell antiviral activity in coculture. Gating strategy to quantify the inhibition of Caco2 cells infection in the presence of MK-8722 and/or CD8^+^ T cells from vaccinated individuals shown in Figure 5C. Infection was evaluated within Caco2 cells corresponding to the CD3^-^CD8^-^ population. The low number of CD8^+^ T cells (around 150) prevented us from investigating their activation level.

## References

1. Fernandes, Q. et al. Emerging COVID-19 variants and their impact on SARS-CoV-2 diagnosis, therapeutics and vaccines. Ann Med 54, 524–540 (2022).

2. Haque, Ashwaq, O., Sarief, A. & Azad John Mohamed, A. K. A comprehensive review about SARS-CoV-2. Future Virol 15, 625–648 (2020).

3. Ji, M. et al. VMP1 and TMEM41B are essential for DMV formation during β-coronavirus infection. J Cell Biol 221, e202112081 (2022).

4. Snijder, E. J. et al. A unifying structural and functional model of the coronavirus replication organelle: Tracking down RNA synthesis. PLOS Biology 18, e3000715 (2020).

5. Farley, S. E. et al. A global lipid map reveals host dependency factors conserved across SARS-CoV-2 variants. Nat Commun 13, 3487 (2022).

6. Tabata, K. et al. Convergent use of phosphatidic acid for hepatitis C virus and SARS-CoV-2 replication organelle formation. Nat Commun 12, 7276 (2021).

7. Tanner, J. E. & Alfieri, C. The Fatty Acid Lipid Metabolism Nexus in COVID-19. Viruses 13, 90 (2021).

8. Bajimaya, S., Frankl, T., Hayashi, T. & Takimoto, T. Cholesterol is required for stability and infectivity of influenza A and respiratory syncytial viruses. Virology 510, 234–241 (2017).

9. Cheng, N. et al. Protein post-translational modification in SARS-CoV-2 and host interaction. Front Immunol 13, 1068449 (2022).

10. McBride, C. E. & Machamer, C. E. Palmitoylation of SARS-CoV S protein is necessary for partitioning into detergent-resistant membranes and cell–cell fusion but not interaction with M protein. Virology 405, 139–148 (2010).

11. Mesquita, F. S. et al. S-acylation controls SARS-CoV-2 membrane lipid organization and enhances infectivity. Dev Cell 56, 2790–2807.e8 (2021).

12. Dias, S. S. G. et al. Lipid droplets fuel SARS-CoV-2 replication and production of inflammatory mediators. PLoS Pathog 16, e1009127 (2020).

13. Grootemaat, A. E. et al. Lipid and Nucleocapsid N-Protein Accumulation in COVID-19 Patient Lung and Infected Cells. Microbiology Spectrum 10, e01271–21 (2022).

14. Nardacci, R. et al. Evidences for lipid involvement in SARS-CoV-2 cytopathogenesis. Cell Death Dis 12, 1–12 (2021).

15. Wang, W. et al. Genetic variety of ORF3a shapes SARS-CoV-2 fitness through modulation of lipid droplet. J Med Virol 95, e28630 (2023).

16. Chu, J. et al. Pharmacological inhibition of fatty acid synthesis blocks SARS-CoV-2 replication. Nat Metab 3, 1466–1475 (2021).

17. Koepke, L., Hirschenberger, M., Hayn, M., Kirchhoff, F. & Sparrer, K. M. Manipulation of autophagy by SARS-CoV-2 proteins. Autophagy 17, 2659–2661 (2021).

18. Gassen, N. C. et al. SARS-CoV-2-mediated dysregulation of metabolism and autophagy uncovers host-targeting antivirals. Nat Commun 12, 3818 (2021).

19. Zhou, H., Hu, Z. & Castro-Gonzalez, S. Bidirectional interplay between SARS-CoV-2 and autophagy. mBio 0, e01020–23 (2023).

20. Zhang, Y., Liu, S., Xu, Q., Li, H. & Lu, K. Cleavage of the selective autophagy receptor SQSTM1/p62 by the SARS-CoV-2 main protease NSP5 prevents the autophagic degradation of viral membrane proteins. Mol Biomed 3, 17 (2022).

21. Twu, W.-I. et al. Contribution of autophagy machinery factors to HCV and SARS-CoV-2 replication organelle formation. Cell Rep 37, 110049 (2021).

22. Hou, P. et al. The ORF7a protein of SARS-CoV-2 initiates autophagy and limits autophagosome-lysosome fusion via degradation of SNAP29 to promote virus replication. Autophagy 19, 551–569 (2023).

23. Koyama-Honda, I., Itakura, E., Fujiwara, T. K. & Mizushima, N. Temporal analysis of recruitment of mammalian ATG proteins to the autophagosome formation site. Autophagy 9, 1491–1499 (2013).

24. Miao, G. et al. ORF3a of the COVID-19 virus SARS-CoV-2 blocks HOPS complex-mediated assembly of the SNARE complex required for autolysosome formation. Dev Cell 56, 427–442.e5 (2021).

25. Zhang et al. The SARS-CoV-2 protein ORF3a inhibits fusion of autophagosomes with lysosomes. Cell Discov 7, 31 (2021).

26. Bizzotto, J. et al. SARS-CoV-2 Infection Boosts MX1 Antiviral Effector in COVID-19 Patients. iScience 23, 101585 (2020).

27. Danziger, O., Patel, R. S., DeGrace, E. J., Rosen, M. R. & Rosenberg, B. R. Inducible CRISPR activation screen for interferon-stimulated genes identifies OAS1 as a SARS-CoV-2 restriction factor. PLOS Pathogens 18, e1010464 (2022).

28. Li, X. et al. SARS-CoV-2 ORF10 suppresses the antiviral innate immune response by degrading MAVS through mitophagy. Cell Mol Immunol 19, 67–78 (2022).

29. Smith, N. et al. Defective activation and regulation of type I interferon immunity is associated with increasing COVID-19 severity. Nat Commun 13, 7254 (2022).

30. Øynebråten, I. Involvement of autophagy in MHC class I antigen presentation. Scand J Immunol 92, e12978 (2020).

31. Wen, Z. et al. Enhancement of SARS-CoV-2 N Antigen-Specific T Cell Functionality by Modulating the Autophagy-Mediated Signal Pathway in Mice. Viruses 15, 1316 (2023).

32. Zhang et al. The ORF8 protein of SARS-CoV-2 mediates immune evasion through down-regulating MHC-Ι. Proc Natl Acad Sci U S A 118, e2024202118 (2021).

33. Steinberg, G. R. & Hardie, D. G. New insights into activation and function of the AMPK. Nat Rev Mol Cell Biol 24, 255–272 (2023).

34. Moreira, D. et al. AMP-activated protein kinase as a target for pathogens: friends or foes? Curr Drug Targets 17, 942–953 (2016).

35. Farfan-Morales, C. N. et al. The antiviral effect of metformin on zika and dengue virus infection. Sci Rep 11, 8743 (2021).

36. Myers, R. W. et al. Systemic pan-AMPK activator MK-8722 improves glucose homeostasis but induces cardiac hypertrophy. Science 357, 507–511 (2017).

37. Muise, E. S. et al. Pharmacological AMPK activation induces transcriptional responses congruent to exercise in skeletal and cardiac muscle, adipose tissues and liver. PLOS ONE 14, e0211568 (2019).

38. Cottignies-Calamarte, A., He, F., Zhu, A., Real, F. & Bomsel, M. Protocol to detect infectious SARS-CoV-2 at low levels using in situ hybridization techniques. STAR Protocols 4, 102593 (2023).

39. Desmarets, L. et al. A reporter cell line for the automated quantification of SARS-CoV-2 infection in living cells. Front Microbiol 13, 1031204 (2022).

40. Lu, X. et al. US CDC Real-Time Reverse Transcription PCR Panel for Detection of Severe Acute Respiratory Syndrome Coronavirus 2. Emerg Infect Dis 26, 1654–1665 (2020).

41. Decker, T. The early interferon catches the SARS-CoV-2. Journal of Experimental Medicine 218, e20211667 (2021).

42. Zhou, Q. et al. Interferon-α2b Treatment for COVID-19. Front Immunol 11, 1061 (2020).

43. Schroeder, S. et al. Interferon antagonism by SARS-CoV-2: a functional study using reverse genetics. The Lancet Microbe 2, e210–e218 (2021).

44. Liu, J., et al. CD8 T cells contribute to vaccine protection against SARS-CoV-2 in macaques. Sci Immunol 7, eabq7647 (2022).

45. Taus, E. et al. Dominant CD8+ T Cell Nucleocapsid Targeting in SARS-CoV-2 Infection and Broad Spike Targeting From Vaccination. Front Immunol 13, 835830 (2022).

46. Vitiello, L. et al. Long Lasting Cellular Immune Response Induced by mRNA Vaccination: Implication for Prevention Strategies. Front Immunol 13, 836495 (2022).

47. Parthasarathy, H., Tandel, D., Siddiqui, A. H. & Harshan, K. H. Metformin suppresses SARS-CoV-2 in cell culture. Virus Research 323, 199010 (2023).

48. Bonnet, R., Mariault, L. & Peyron, J.-F. Identification of potentially anti-COVID-19 active drugs using the connectivity MAP. PLoS One 17, e0262751 (2022).

49. Doshi, H. et al. AMPK protects endothelial cells against HSV-1 replication via inhibition of mTORC1 and ACC1. Microbiol Spectr 11, e0041723 (2023).

50. Farfan-Morales, C. N. et al. Anti-flavivirus Properties of Lipid-Lowering Drugs. Frontiers in Physiology 12, (2021).

51. Jiménez de Oya, N., Blázquez, A.-B., Casas, J., Saiz, J.-C. & Martín-Acebes, M. A. Direct Activation of Adenosine Monophosphate-Activated Protein Kinase (AMPK) by PF-06409577 Inhibits Flavivirus Infection through Modification of Host Cell Lipid Metabolism. Antimicrob Agents Chemother 62, e00360–18 (2018).

52. Prasad, V. et al. Enhanced SARS-CoV-2 entry via UPR-dependent AMPK-related kinase NUAK2. Molecular Cell 83, 2559–2577.e8 (2023).

53. Yuan, W.-C. et al. NUAK2 is a critical YAP target in liver cancer. Nat Commun 9, 4834 (2018).

54. Erickson, S. M. et al. Metformin for Treatment of Acute COVID-19: Systematic Review of Clinical Trial Data Against SARS-CoV-2. Diabetes Care 46, 1432–1442 (2023).

55. Foretz, M., Even, P. C. & Viollet, B. AMPK Activation Reduces Hepatic Lipid Content by Increasing Fat Oxidation In Vivo. Int J Mol Sci 19, E2826 (2018).

56. Yeudall, S. et al. Macrophage acetyl-CoA carboxylase regulates acute inflammation through control of glucose and lipid metabolism. Sci Adv 8, eabq1984 (2022).

57. Fomin, G. et al. Cytokine response and damages in the lungs of aging Syrian hamsters on a high-fat diet infected with the SARS-CoV-2 virus. Front Immunol 14, 1223086 (2023).

58. Tsai, P.-H. et al. Modifications of lipid pathways restrict SARS-CoV-2 propagation in human induced pluripotent stem cell-derived 3D airway organoids. Journal of Advanced Research (2023) doi:10.1016/j.jare.2023.08.005.

59. Yuan, Y. et al. The CH24H metabolite, 24HC, blocks viral entry by disrupting intracellular cholesterol homeostasis. Redox Biol 64, 102769 (2023).

60. Santosa, A. et al. Protective effects of statins on COVID-19 risk, severity and fatal outcome: a nationwide Swedish cohort study. Sci Rep 12, 12047 (2022).

61. Teixeira, L. et al. Simvastatin Downregulates the SARS-CoV-2-Induced Inflammatory Response and Impairs Viral Infection Through Disruption of Lipid Rafts. Front Immunol 13, 820131 (2022).

62. Paunovic, V. et al. Autophagy Receptor p62 Regulates SARS-CoV-2-Induced Inflammation in COVID-19. Cells 12, 1282 (2023).

63. Tan, X. et al. Coronavirus subverts ER-phagy by hijacking FAM134B and ATL3 into p62 condensates to facilitate viral replication. Cell Rep 42, 112286 (2023).

64. Thinwa, J. W. et al. CDKL5 regulates p62-mediated selective autophagy and confers protection against neurotropic viruses. J Clin Invest 134, e168544 (2024).

65. Dong, X. et al. Sorting nexin 5 mediates virus-induced autophagy and immunity. Nature 589, 456–461 (2021).

66. Gassen, N. C. et al. SKP2 attenuates autophagy through Beclin1-ubiquitination and its inhibition reduces MERS-Coronavirus infection. Nat Commun 10, 5770 (2019).

67. Kumar, A. V., Mills, J. & Lapierre, L. R. Selective Autophagy Receptor p62/SQSTM1, a Pivotal Player in Stress and Aging. Frontiers in Cell and Developmental Biology 10, (2022).

68. Gestal-Mato, U. & Herhaus, L. Autophagy-dependent regulation of MHC-I molecule presentation. J Cell Biochem (2023) doi:10.1002/jcb.30416.

69. Manzoni, T. B. & López, C. B. Defective (interfering) viral genomes re-explored: impact on antiviral immunity and virus persistence. Future Virol 13, 493–503 (2018).

70. Yang, E. & Li, M. M. H. All About the RNA: Interferon-Stimulated Genes That Interfere With Viral RNA Processes. Front Immunol 11, 605024 (2020).

71. Abe, K. et al. Analysis of interferon-beta mRNA stability control after poly(I:C) stimulation using RNA metabolic labeling by ethynyluridine. Biochem Biophys Res Commun 428, 44–49 (2012).

72. Cabral-Piccin, M. P. et al. Primary role of type I interferons for the induction of functionally optimal antigen-specific CD8+ T cells in HIV infection. EBioMedicine 91, 104557 (2023).

73. Hemann, E. A. et al. A Small Molecule RIG-I Agonist Serves as an Adjuvant to Induce Broad Multifaceted Influenza Virus Vaccine Immunity. J Immunol 210, 1247–1256 (2023).

74. Proietti, E. et al. Type I IFN as a Natural Adjuvant for a Protective Immune Response: Lessons from the Influenza Vaccine Model1. The Journal of Immunology 169, 375–383 (2002).

75. Sasaki, E. et al. Systemically inoculated adjuvants stimulate pDC-dependent IgA response in local site. Mucosal Immunol 16, 275–286 (2023).

